# Dual targeting factors are required for LXG toxin export by the bacterial type VIIb secretion system

**DOI:** 10.1101/2022.07.06.499029

**Authors:** Timothy A. Klein, Dirk W. Grebenc, Prakhar Y. Shah, Owen D. McArthur, Brandon H. Dickson, Michael G. Surette, Youngchang Kim, John C. Whitney

**Affiliations:** Michael DeGroote Institute for Infectious Disease Research, McMaster University, Hamilton, ON, L8S 4K1, Canada; Department of Biochemistry and Biomedical Sciences, McMaster University, Hamilton, ON, L8S 4K1, Canada; Department of Medicine, Farncombe Family Digestive Health Research Institute, McMaster University, Hamilton, ON, L8S 4K1, Canada; Structural Biology Center, X-ray Science Division, Advanced Photon Source, Argonne National Laboratory, Lemont, Illinois, 60439, USA; David Braley Centre for Antibiotic Discovery, McMaster University, Hamilton, ON, L8S 4K1, Canada

**Author notes:** To whom correspondence should be addressed: J.C.W.

**Keywords:** Type VII secretion system, protein secretion, protein-protein interactions, bacterial antagonism, antibacterial toxins, X-ray crystallography

## Abstract

Bacterial type VIIb secretion systems (T7SSb) are multi-subunit integral membrane protein complexes found in Firmicutes that play a role in both bacterial competition and virulence by secreting toxic effector proteins. The majority of characterized T7SSb effectors adopt a polymorphic domain architecture consisting of a conserved N-terminal Leu-X-Gly (LXG) domain and a variable C-terminal toxin domain. Recent work has started to reveal the diversity of toxic activities exhibited by LXG effectors; however, little is known about how these proteins are recruited to the T7SSb apparatus. In this work, we sought to characterize genes encoding domains of unknown function (DUFs) 3130 and 3958, which frequently co-occur with LXG effector-encoding genes. Using coimmunoprecipitation-mass spectrometry analyses, *in vitro* copurification experiments and T7SSb secretion assays, we find that representative members of these protein families form heteromeric complexes with their cognate LXG domain and in doing so, function as targeting factors that promote effector export. Additionally, an X-ray crystal structure of a representative DUF3958 protein, combined with predictive modelling of DUF3130 using AlphaFold2, reveals structural similarity between these protein families and the ubiquitous WXG100 family of T7SS effectors. Interestingly, we identify a conserved FxxxD motif within DUF3130 that is reminiscent of the YxxxD/E “export arm” found in Mycobacterial T7SSa substrates and mutation of this motif abrogates LXG effector secretion. Overall, our data experimentally link previously uncharacterized bacterial DUFs to type VIIb secretion and reveal a molecular signature required for LXG effector export.

**Significance statement:** Type VIIb secretion systems (T7SSb) are protein secretion machines used by an array of Gram-positive bacterial genera including *Staphylococcus, Streptococcus, Bacillus*, and *Enterococcus*. These bacteria use the T7SSb to facilitate interbacterial killing and pathogenesis through the secretion of toxins. Although the modes of toxicity for a number of these toxins have been investigated, the mechanisms by which they are recognized and secreted by T7SSb remains poorly understood. The significance of this work is the discovery of two new protein families, termed Lap1 and Lap2, that directly interact with these toxins and are required for their secretion. Overall, Lap1 and Lap2 represent two widespread families of proteins that function as targeting factors that participate in T7SSb-dependent toxin release from Gram-positive bacteria.

## INTRODUCTION

Protein secretion is an essential aspect of bacterial physiology that plays a critical role in diverse cellular activities including interbacterial competition and infection of host cells (1, 2). Bacteria possess several protein export pathways, often referred to as secretion systems, that facilitate protein transport across the cell envelope. In general, these pathways consist of membrane proteins that form the secretion apparatus and effector proteins that transit the secretion system. One important property of protein secretion apparatuses is their ability to recognize and export a specific set of effector proteins among the myriad cytosolic proteins within a cell. In many well-characterized examples, effectors harbour a signal sequence that is recognized by the secretion apparatus and the recruitment of this signal sequence to the apparatus often requires the involvement of effector-specific accessory proteins (3–7).

Bacteria encode two ubiquitous secretion systems known as the general secretory pathway (Sec) and the Twin-arginine translocase (TAT). In addition to Sec and TAT, many Gram-negative bacteria encode a series of specialized secretion systems, several of which span the entirety of the diderm cell envelope (2). By contrast, a substantial number of Gram-positive bacteria possess a single specialized secretion pathway referred to as the type VII secretion system (T7SS)(8). In recent years, this pathway has been further differentiated into two subtypes, T7SSa and T7SSb, to reflect the substantial differences in protein subunits that comprise each secretion apparatus (9). The T7SSa is found in Actinobacteria where it functions as an essential virulence factor for many pathogenic species of Mycobacteria with specific T7SSa pathways linked to diverse functions including phagosomal escape, metal ion homeostasis, and conjugation (10–13). The T7SSb is found in Firmicutes and is involved in the pathogenesis of *Staphylococcus aureus, Streptococcus agalactiae* and *Streptococcus intermedius* (14–16). In addition, several recent studies have uncovered a role for this pathway in mediating antagonistic interbacterial interactions in *S. aureus, S. intermedius, Enterococcus faecalis* and *Bacillus subtilis* (17–20). Both T7SS subtypes have an FtsK-SpoIIIE family ATPase known as EccC/EssC that is thought to energize effector secretion and export one or more small α-helical effectors belonging to the WXG100 protein family (8). Beyond these similarities, T7SSa and T7SSb require different sets of apparatus proteins and export different families of effector proteins (9).

LXG proteins are emerging as the predominant group of effectors exported by T7SSb pathways (18–21). These proteins possess a polymorphic domain architecture comprised of a conserved ~200 amino acid N-terminal LXG (Leu-X-Gly) domain and a variable C-terminal toxin domain (22). The toxic activities of several LXG effectors have been biochemically characterized and includes toxin domains that hydrolyze NAD^+^, disrupt peptidoglycan biosynthesis, depolarize membranes, and degrade essential nucleic acids (17, 18, 21, 23). By contrast, little is known about the function of LXG domains. Based on comparisons to other polymorphic toxin systems, Zhang and Aravind propose a role for this domain in effector recruitment to the T7SS apparatus (22). This hypothesis is bolstered by recent bacterial two-hybrid analyses showing that an LXG domain encoded by *B. subtilis* physically interacts with the T7SSb subunit YukC/EssB (24). However, because this experiment relied on a heterologous expression system, it remains unclear if this interaction is sufficient to promote effector secretion or if other factors are additionally required. In support of the need for additional secretion factors, the three LXG effectors exported by *S. intermedius* B196 interact with effector specific Wxg proteins via their LXG domains. Furthermore, it was shown for the TelC effector that its cognate Wxg protein, WxgC, is required for its export (18).

In contrast to LXG proteins, the secretion determinants for T7SSa effectors are better defined. In general, T7SSa effectors exist as obligate heterodimers and heterodimerization is a prerequisite for secretion. The archetypal examples are EsxA (ESAT-6) and EsxB (CFP-10), which are secreted as a heterodimer by the ESX-1 system of *Mycobacterium tuberculosis* (25, 26). The co-secretion of these effectors requires a conserved YxxxD/E motif present at the unstructured C-terminus of EsxB (27). Biochemical characterization of the interaction between the EccC motor ATPase and EsxB suggests that this secretion signal facilitates effector export by inducing EccC multimerization (28). Like EsxB, other families of T7SSa effectors such as EspB and members of the proline-glutamate (PE)/proline-proline-glutamate (PPE) family possess YxxxD/E motifs and in all tested cases, this motif is required for effector export (29–31).

In the present study, we sought to systematically characterize the secretion determinants of LXG effectors exported by the T7SSb pathway. Using two model effectors from two different strains of *S. intermedius*, we find that members of the DUF3130 (also known as TIGR04197, and “Type VII secretion effector, SACOL2603 family”) and the DUF3958 protein families function as dual targeting factors that physically interact with and promote the secretion of their cognate effector. Using structural analyses, we find that DUF3130 and DUF3958 bear resemblance to WXG100 effectors; however, in contrast to these effectors, they are not exported by the T7SSb. While DUF3958 proteins lack conserved sequence motifs that could provide insight into their precise function, DUF3130 proteins possess a highly conserved FxxxD motif that resembles the secretion signal found in T7SSa substrates. Moreover, site-specific mutation of this motif abrogates LXG effector export. Overall, our work uncovers new intracellular factors involved in LXG effector secretion, provides molecular insights into how these factors function, and demonstrates that effector secretion by T7SSb pathways may share more similarities to their Mycobacterial T7SSa counterparts than previously appreciated.

## RESULTS

### A DUF3958 protein is required for export of the LXG effector TelC from *S. intermedius* B196

In a recent bioinformatics study on the LXG effector repertoire of *Listeria monocytogenes*, Bowran and Palmer noted the near ubiquitous existence of two small open reading frames upstream of LXG genes (32). In our initial characterization of the model LXG effector TelC from *S. intermedius* B196 we found that the protein product of one of these genes, WxgC, physically interacts with TelC and is required for its T7SSb-dependent export (18). However, the function of the other LXG effector associated gene, SIR_1490, was not examined. WxgC and SIR_1490 have homology to the DUF3130 and DUF3958 protein families, respectively. Given the frequent co-occurrence of the genes encoding these proteins within LXG effector gene clusters, we hypothesized that SIR_1490 also plays a role in the export of TelC (Fig 1A). To test this, we first generated an *S. intermedius* B196 strain lacking SIR_1490 and examined the ability of this strain to export TelC into culture supernatants. In line with our hypothesis, only intracellular TelC was detected in ΔSIR_1490 and TelC secretion could be restored by plasmid-borne expression of SIR_1490 (Fig 1B). Export of the WXG100 effector EsxA, a hallmark of a functional T7SS apparatus, was unaffected by mutational inactivation of SIR_1490 indicating that the loss of TelC secretion is not due to a defect in T7SSb apparatus function (Fig 1C)(8). Like EsxA, WxgC and SIR_1490 are predicted to be α-helical proteins of approximately 100 amino acids in length (Fig S1). Therefore, we next considered the possibility that these proteins are also exported by the T7SSb. In contrast to EsxA and TelC, we were unable to detect either of these proteins in the culture medium (Fig 1D). Based on these data, we conclude that WxgC and SIR_1490 function as cytoplasmic factors that facilitate the T7SSb-dependent export of TelC. In light of these and subsequent findings, we propose to rename WxgC/SIR_1491 and name SIR_1490 to <u>L</u>XG-associated <u>α</u>-helical <u>p</u>rotein for Tel<u>C</u> 1 (LapC1, DUF3130) and 2 (LapC2, DUF3958), respectively.

**Figure 1.**
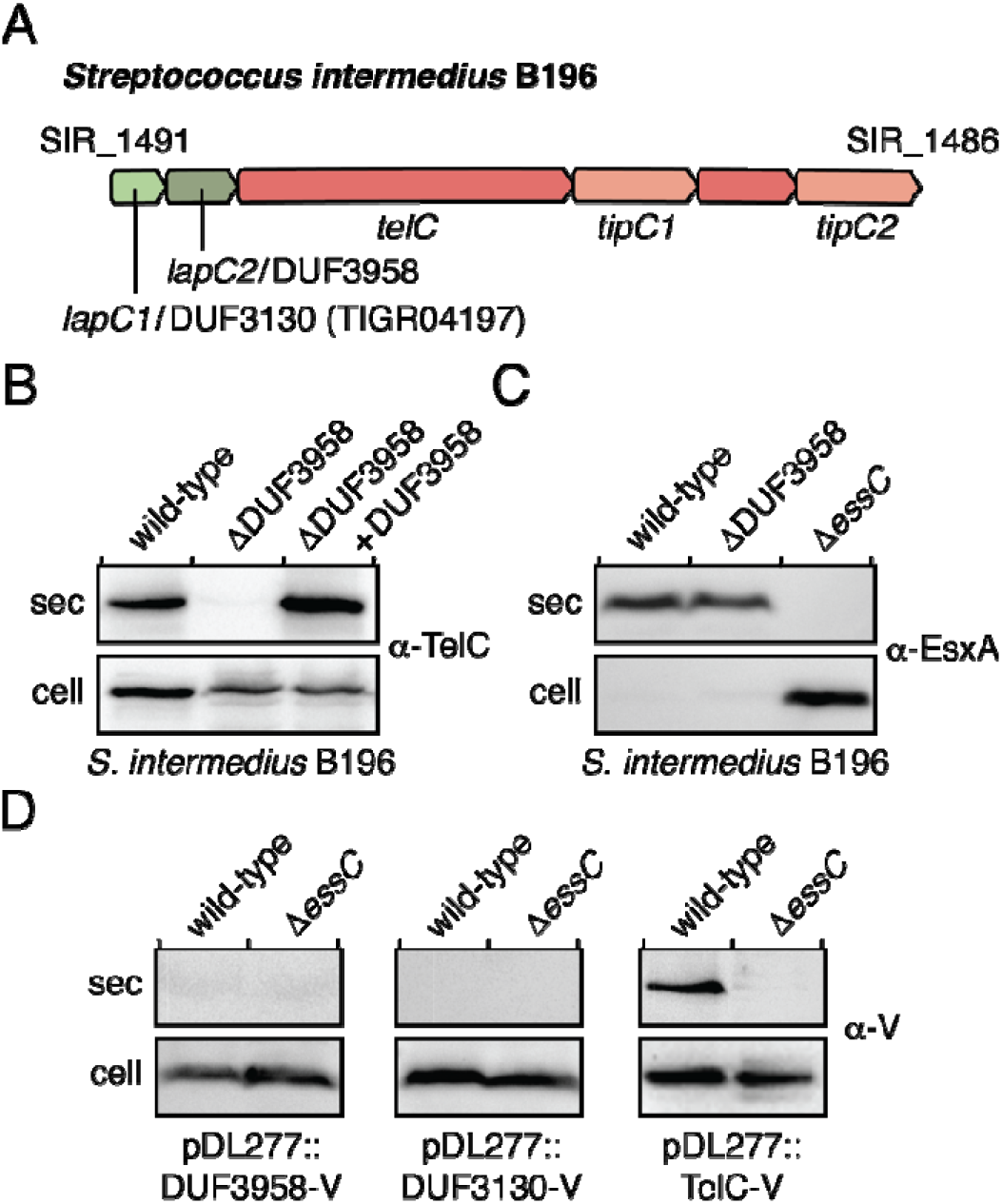
SIR_1490 encodes a DUF3958 protein required for the export of LXG effector TelC. (A) Schematic of the *telC* gene cluster from *S. intermedius* B196. Locus tags and gene names/DUF families are provided above and below the gene diagram, respectively. Genes are coloured to signify their function/context: light green – DUF3130 homolog (*lapC1*), dark green – DUF3958 homolog (*lapC2*), orange – LXG effector (*telC*) or orphan toxin domain (SIR_1487), salmon – immunity genes. (B-D) Western blot analysis of the secreted (sec) and cell fractions of the indicated *S. intermedius* B196 strains. Protein specific antibodies were used to detect endogenous TelC and EsxA (B and C) and anti-VSV-G epitope antibody was used to detect ectopically expressed VSV-G-tagged SIR_1490 (DUF3958-V) and VSV-G-tagged SIR_1491 (DUF3130-V) (D). *S. intermedius* B196 *ΔessC* is a T7SS-deficient control. The pDL277::DUF3958-V complementation vector used in (B) is the same as that used to assess secretion in (D).

### TelC, LapC1 and LapC2 physically interact to form a heterotrimeric pre-secretion complex

Given our genetic data linking both *lapC1* and *lapC2* to the T7SSb-dependent export of TelC, we next wanted to examine whether the encoded proteins physically interact with TelC in the context of their native organism. To probe this, we expressed Vesicular Stomatitis Virus G (VSV-G) epitope tagged TelC (TelC-V) in *S. intermedius* B196, performed an immunoprecipitation using anti-VSV-G antibody, and identified proteins that were enriched relative to a control strain by mass spectrometry (Fig 2A and Supplemental Table S1). LapC1 was highly enriched in the TelC-V expressing strain, corroborating previous bacterial two-hybrid data in *E. coli* that indicated these proteins interact directly (18). Interestingly, LapC2 was also highly enriched, suggesting that LapC2 also interacts with TelC. PepC, an aminopeptidase with no known role in type VII secretion, was also present in our TelC-V sample and absent in our control, although it was present in lower overall abundance as measured by total spectral counts (33). We speculate that a small amount of PepC may interact with highly expressed proteins under some conditions but consider it unlikely that PepC is a bonafide interaction partner of TelC. To substantiate this assumption and to validate TelC’s interaction with LapC1 and LapC2, we next performed a similar immunoprecipitation using VSV-G tagged LapC1 (LapC1-V) as the bait protein (Fig 2B and Supplemental Table S1). In this experiment, both TelC and LapC2 were enriched relative to the control sample, but PepC was not. Based on these data, we conclude that TelC, LapC1 and LapC2 interact to form an effector pre-secretion complex in *S. intermedius* B196.

**Figure 2.**
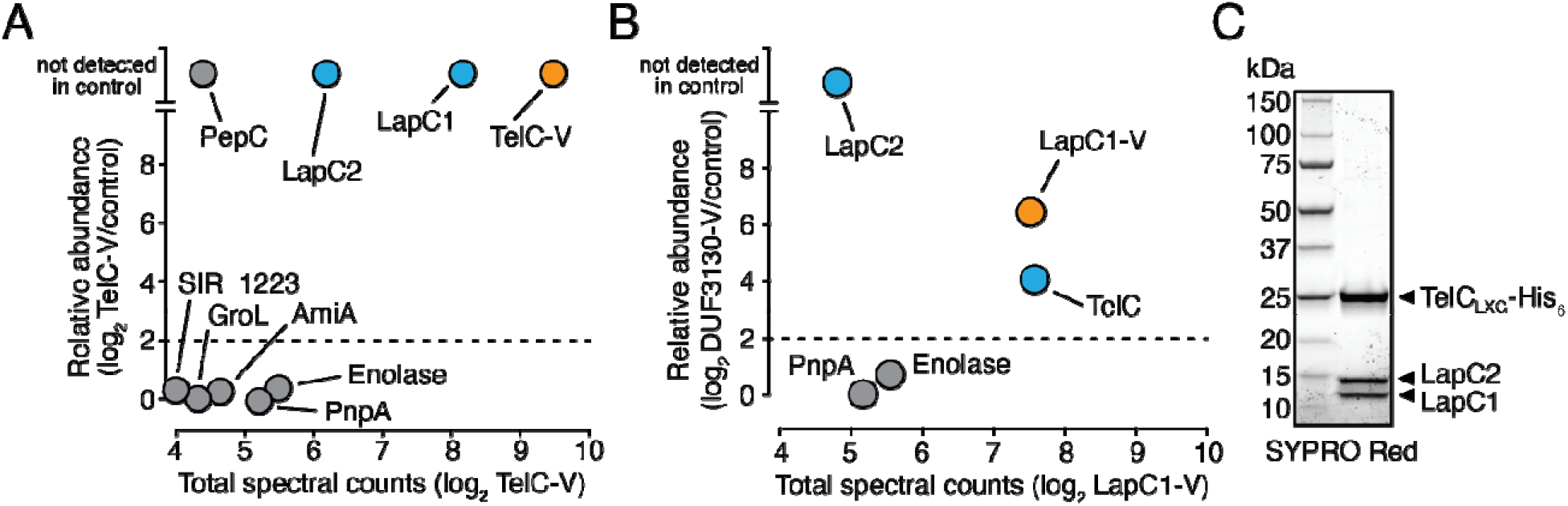
LapC1 and LapC2 interact with the LXG domain of TelC to form an effector pre-secretion complex. (A and B) Mass spectrometry analysis of immunoprecipitated VSV-G tagged TelC (TelC-V) (A) and LapC1 (LapC1-V) (B). Total spectral counts of abundantly detected proteins and fold enrichment relative to a control strain are plotted on the X- and Y-axes, respectively. In both panels, the immunoprecipitated protein is coloured orange while interaction partners are coloured blue. (C) SYPRO Red stained gel showing purified TelC_LXG_–LapC1–LapC2 complex. Proteins were co-expressed in *E. coli* and purified using nickel affinity and size exclusion chromatography.

Because protein-protein interactions identified by co-immunoprecipitation can be indirect in nature, we next attempted to co-express and purify TelC with LapC1 and LapC2 using an *E. coli* overexpression system. Previous bacterial two-hybrid data showed that the LXG domain of TelC (TelC_LXG_) is both necessary and sufficient for LapC1 interaction (18). Therefore, we similarly used TelC_LXG_ to assess TelC-LapC1-LapC2 heteromer formation. Using His_6_-tagged TelC_LXG_ to facilitate nickel affinity chromatography, we found that TelC_LXG_ copurified with both LapC1 and LapC2 after nickel affinity and size exclusion chromatography (Fig 2C). Taken together, our results indicate that the physical association of LapC1 and LapC2 with TelC’s LXG domain promotes TelC export by the T7SS of *S. intermedius* B196.

### TelD is a novel LXG-containing T7SSb effector that also requires a cognate Lap1-Lap2 pair for export

To test the generalizability of our findings on TelC, LapC1 and LapC2, we next sought to determine if a second LXG effector also requires heterocomplex formation with a cognate Lap1-Lap2 pair to facilitate its secretion by the T7SS. To this end, we examined the recently sequenced GC1825 strain of *S. intermedius* and identified a candidate LXG-domain containing T7SS effector, which we named *telD* to remain consistent with the established *S. intermedius* T7SS LXG effector nomenclature (18). The *telD* gene neighbourhood has similar synteny to that of *telC* in that the effector gene is found immediately downstream of *lapC1* and *lapC2* homologous genes, which we henceforth refer to as *lapD1* and *lapD2*, respectively, to reflect their genetic linkage to *telD* (Fig 3A). Downstream of *telD* are two DUF443-encoding genes, which belong to a family of proteins that contain TsaI, a characterized immunity protein for the membrane depolarizing LXG effector TspA of *S. aureus* (21). The final ORF in the predicted operon is a DUF4176-encoding gene, members of which are often found among T7SS genes but whose function is unknown (9, 34).

**Figure 3.**
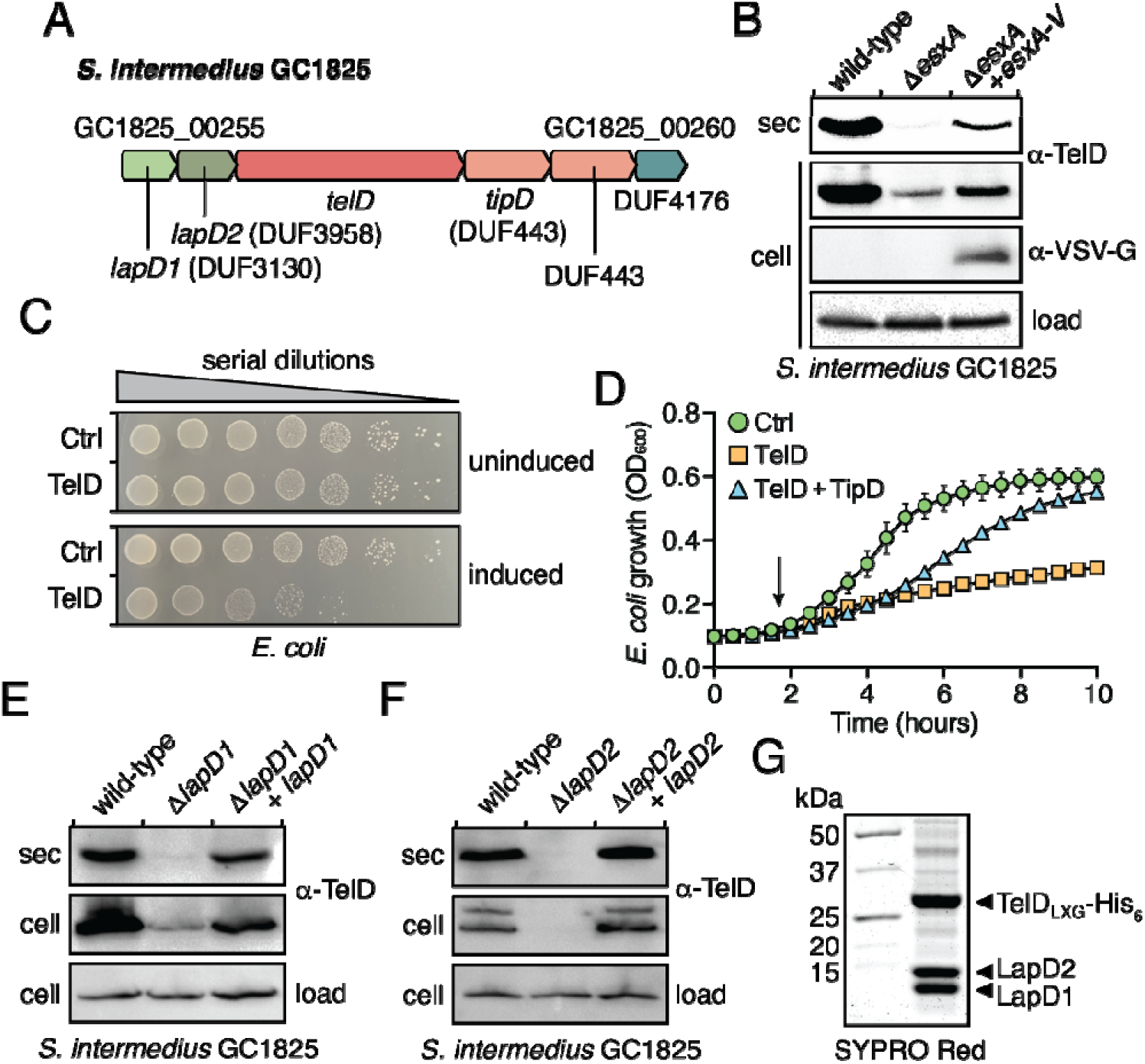
Lap1 and Lap2 proteins are required for secretion of the novel LXG effector TelD. (A) Schematic of the *telD* gene cluster from *S. intermedius* GC1825. Locus tags and gene names/DUF families are provided above and below the gene diagram, respectively. Genes are coloured to signify their function/context: light green – DUF3130 homolog (*lapD1*), dark green – DUF3958 homolog (*lapD2*), orange – LXG effector (*telD*), salmon – immunity genes, blue – DUF4176. (B-D) *S. intermedius* GC1825 TelD is an antibacterial toxin that is exported in a T7SS-dependent manner. Western blot analysis of the secreted (sec) and cell fractions of the indicated *S. intermedius* GC1825 strains (B). CFU plating (C) and growth curves (D) of *E. coli* cells expressing TelD, TelD with the TipD immunity protein or a vector control (Ctrl). In panel D, arrow indicates when inducer was added, and error bars represent SEM. (E and F) Western blot analysis of the secreted and cell fractions of the indicated *S. intermedius* GC1825 strains. (G) SYPRO Red stained gel of purified TelD_LXG_–LapD1–LapD2 complex. In all panels containing western blots, a TelD specific antibody was used to detect endogenous TelD and a cross-reactive band was used as a loading control.

We first wanted to determine if TelD is indeed a T7SS effector as would be predicted due to it possessing an N-terminal LXG domain. To test this, we deleted the gene encoding the essential T7SSb component, *esxA*, and examined TelD secretion by western blot using a TelD-specific antibody. Our results show that in contrast to wild-type *S. intermedius* GC1825, the T7SS-inactivated strain is unable to secrete TelD (Fig 3B). Of note, we observed lower levels of intracellular TelD in the *ΔesxA* strain relative to wild-type and the reason for this is currently unclear. Nonetheless, complementing *esxA* in trans resulted in a partial restoration of TelD export suggesting that TelD is secreted in a T7SS-dependant manner. Antibacterial activity is a property of all LXG toxins characterized to date, so we next wanted to examine if TelD is also toxic to bacterial cells. Consistent with this precedent, we found that expression of the TelD toxin in *E. coli* led to an approximate 100-fold decrease in cell viability (Fig 3C). Furthermore, when grown in liquid culture, we observed that TelD caused *E. coli* growth arrest shortly after induction of toxin expression but did not cause cell lysis (Fig 3D). Finally, co-expression of the adjacent DUF443-encoding gene, henceforth referred to as *tipD*, substantially restored *E. coli* growth (Fig 3D). Given that it shares the same family of predicted immunity proteins as TspA, TelD may similarly inhibit bacterial growth via membrane depolarization. However, while their LXG domains possess 29.4% sequence identity and are predicted to have nearly identical secondary structure, the toxin domains are only 13% identical and yield substantially different structural predictions (Fig S2). Therefore, this putative activity will require experimental validation. In sum, these data indicate that TelD is a T7SS effector with antibacterial properties.

Having established that TelD is a substrate of *S. intermedius* GC1825’s T7SS, we next examined the dependency of its secretion on *lapD1* and *lapD2*. To this end, we generated *S. intermedius* GC1825 strains lacking either *lapD1* or *lapD2*. Interestingly, and in contrast to TelC, we found that overall TelD levels were greatly diminished in the absence of *lapD1* and below the limit of detection in a *lapD2* deletion strain (Fig 3E and 3F). Consistent with our findings on TelC, TelD export was also abrogated in the strain lacking *lapD1*. Importantly, cellular levels of TelD as well as its export via the T7SS could be restored by complementing each deletion strain with a plasmid-borne copy of the deleted gene (Fig 3E and 3F). The decrease in cellular TelD levels differs from our findings with TelC and suggests that LXG effectors have differing levels of intrinsic stability. In the case of TelD, our data indicate that in addition to being required for effector export, LapD1 and LapD2 are exhibiting chaperone-like properties by stabilizing their cognate effector prior to its export from the cell. Similar to TelC, we found that the LXG domain of TelD (TelD_LXG_) forms a stable heteromeric complex with LapD1 and LapD2 when overexpressed in *E. coli* and copurified using nickel affinity and size exclusion chromatography (Fig 3G). In summary, our TelD data corroborate our findings on TelC by showing that LXG effector secretion, and in some cases LXG effector stability, requires the activities of genetically linked Lap1 and Lap2 proteins.

### A crystal structure of LapD2 reveals its structural similarity to the WXG100 family of T7SS effectors

To better understand the molecular basis for Lap1 and Lap2 function, we initiated protein crystallization experiments on six representative members of each protein family including those linked to TelC and TelD export. Unfortunately, most of the Lap1 and Lap2 proteins that we tested expressed poorly or were recalcitrant to crystallization. Despite this discouraging trend, LapD2 was the sole exception and after systematic optimization of its crystallization conditions it formed crystals that diffracted to 2.2Å. The crystallographic phase problem was overcome using selenomethionine-incorporated protein and the <u>s</u>ingle wavelength <u>a</u>nomalous <u>d</u>ispersion (SAD) technique. The final model of LapD2 was refined to an R_work_/R_free_ of 0.23/0.26 using the native diffraction data (Table 1).

**Table 1.** X-ray data collection and refinement statistics.

|  | LapD2 (selenomethionine) | LapD2 (native) |
| --- | --- | --- |
| **Data Collection** |  |  |
| Wavelength (Å) | 0.9793 | 0.9793 |
| Space group | P3<sub>1</sub>21 | P3<sub>1</sub> |
| Cell dimensions |  |  |
| *a*, *b*, *c* (Å) | 45.2, 45.2, 298.4 | 45.6, 45.6, 298.4 |
| Resolution<sup>a</sup> (Å) | 39.20 – 2.42 (2.46 – 2.42) | 39.46 – 2.20 (2.24 – 2.20) |
| Unique reflections | 14603 (727) | 34956 (1569) |
| CC<sub>1/2</sub><sup>c</sup> | 0.999 (0.371) | 0.999 (0.474) |
| *R*<sub>merge</sub><sup>b</sup> | 0.128 (4.215) | 0.078 (2.118) |
| *R*<sub>pim</sub><sup>c</sup> | 0.036 (1.110) | 0.035 (0.840) |
| *I*/σ*I* | 13.0 (0.6) | 12.9 (0.8) |
| Completeness (%) | 100 (100) | 99.3 (90.6) |
| Redundancy | 21.7 (15.0) | 11.1 (6.6) |
| **Refinement** |  |  |
| *R*<sub>work</sub><sup>d</sup>/ *R*<sub>free</sub> (%) | - | 23.3/26.7 |
| Average B-factors (Å<sup>2</sup>) |  | 76.2 |
| Protein | - | 76.3 |
| Water/Other | - | 56.1/88.5 |
| No. atoms |  |  |
| Protein | - | 3758 |
| Water/Other | - | 35/20 |
| Rms deviations |  |  |
| Bond lengths (Å) | - | 0.005 |
| Bond angles (°) | - | 0.733 |
| Ramachandran plot (%) |  |  |
| Total favored<sup>e</sup> | - | 98.38 |
| Total allowed | - | 1.39 |
| PDB code | - | 7UH4 |

<sup>a</sup>Values in parentheses correspond to the highest resolution shell. <sup>b</sup> $R_{\text{merge}} = \sum_h \sum_j |I_{hj} - \langle I_h \rangle| / \sum_h \sum_j I_{hj}$ , where *I*<sub>hj</sub> is the intensity of observation *j* of reflection *h*. <sup>c</sup>As defined by Karplus and Diederichs (76). <sup>d</sup> $R = \sum_h |F_o| - |F_c| / \sum_h |F_o|$ for all reflections, where *F*<sub>o</sub> and *F*<sub>c</sub> are observed and calculated structure factors, respectively. *R*<sub>free</sub> is calculated analogously for the test reflections, randomly selected and excluded from the refinement. <sup>e</sup>As defined by Molprobit (77)

The overall structure of LapD2 shows that it adopts a helix-turn-helix fold that is reminiscent of the WXG100 family of small secreted T7SS effectors (Fig 4A). However, in contrast to characterized WXG100 proteins, which typically form head-to-toe homodimers mediated by hydrophobic interactions, the turn region of LapD2 contains an intermolecular disulfide bond formed by cysteine 59 that facilitates head-to-head dimerization (Fig 4A and 4B)(35). The head region of LapD2 also possesses a hydrophobic patch that may also contribute to dimerization (Fig 4C). Not surprisingly, the energy of head-to-head dimer formation as predicted by the PDBePISA webtool is highly favourable (Δ^i^G = −21.0) due to the combined effects of burying a hydrophobic patch from the aqueous milieu and possessing a disulfide linkage (36). PDBePISA also revealed a toe-to-toe homodimer interface and this interaction was also suggested to be favourable, although with a lower energy of formation (Δ^i^G = −11.6) (Fig S3). A search for proteins that are structurally homologous to LapD2 using the DALI webserver identified over 12,000 proteins with significant similarity (Z-score > 2) (37). The enormity of this list is due to helix-turn-helix motifs being a common structural element found in numerous proteins of diverse function with the most frequently occurring in our list being DNA-binding proteins. As alluded to above, WXG100 proteins were also well represented with 65 WXG100 family protein structures scoring as significantly similar to LapD2. The top WXG100 hit was a structure of the EsxB protein exported by the ESX-1 T7SSa of *M. tuberculosis* (PDB: 3FAV, Z-score = 8.4, R.M.S.D. = 2.6Å over 90 aligned residues) (Fig S3).

**Figure 4.**
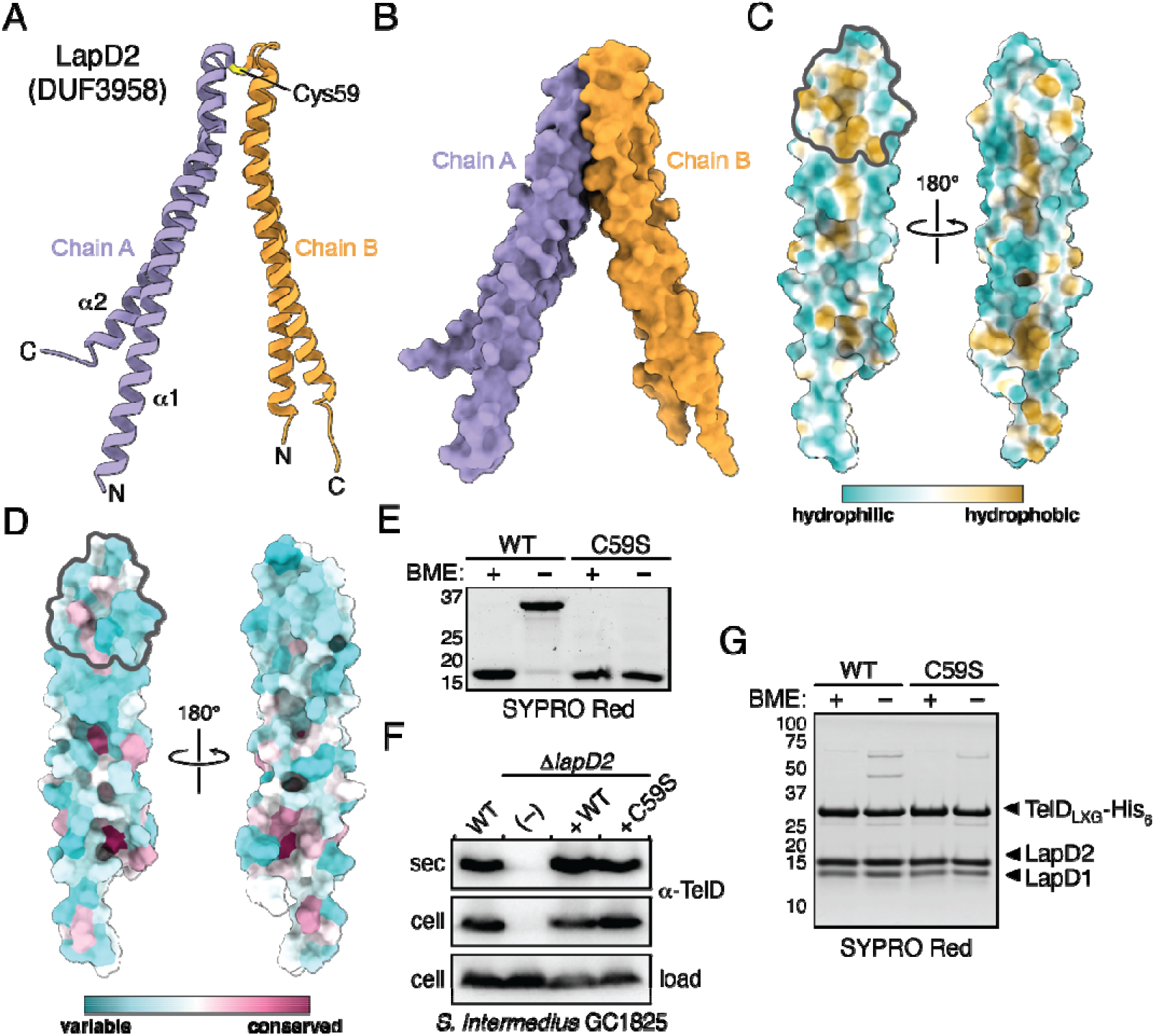
LapD2 is a small α-helical protein reminiscent of WXG100 superfamily proteins. (A and B) Overall structure of LapD2. LapD2 is shown as ribbon (A) and space-filling (B) representations with secondary structure elements, intermolecular disulfide bond, chain identities, and termini labelled where appropriate. (C) Hydrophobicity analysis of LapD2’s surface as calculated by ChimeraX (66). The LapD2 homodimerization interface is denoted by a grey outline. (D) Surface representation of Lap2 sequence conservation mapped onto the LapD2 structure. Sequences used for conservation analysis are available in Supplemental Table S2A. (E-G) Mutation of Cys59 to serine abrogates covalent dimer formation but does not impede TelD secretion or its ability to interact with LapD1 and LapD2. SYPRO Red staining of purified LapD2 and LapD2^C59S^ in the presence or absence of β-mercaptoethanol (BME) (E). Western blot analysis of the secreted and cell fractions of the indicated *S. intermedius* GC1825 strains (F). SYPRO Red staining of purified TelD_LXG_-LapD1-LapD2 and TelD_LXG_-LapD1-LapD2^C59S^ complexes in the presence or absence of BME (G).

We next wanted to determine what structural aspects of LapD2 play a role in facilitating TelD secretion. To initiate this, we first generated a sequence alignment of 95 unique homologous proteins identified using three iterations of the JackHMMER algorithm and mapped the resulting sequence conservation onto the structure of LapD2 (Fig 4D and Supplemental Table S2A). This analysis revealed that Lap2 proteins generally have low sequence conservation. For example, four randomly selected sequences from our list each share approximately 19% pairwise sequence identity to either LapC2 or LapD2 (Fig S4). An alignment using all identified homologs reveals a pattern of hydrophobic residues, particularly leucine, at conserved positions that are interspersed between regions that favour charged and polar residues (Fig S4). Notably, conservation is very low within the interhelical turn region, which contrasts with the highly conserved WXG motif found within structurally similar WXG100 proteins (35). We therefore speculate that shape and/or the surface properties of this protein family may be more critical to function than specific motifs within the primary sequence.

One of the more striking features of LapD2 is the disulfide bond formed by Cys59 that contributes to dimerization. This residue is not conserved among Lap2 proteins indicating that an intermolecular disulfide bond is likely not a universal property of this protein family. Nonetheless, we reasoned that its unique involvement in LapD2 dimerization warranted its functional interrogation in the context of TelD stability and secretion. To accomplish this, we first mutagenized Cys59 to serine (C59S) and confirmed that this variant could no longer form β-mercaptoethanol sensitive dimers *in vitro* (Fig 4E). We next assessed the ability of a strain expressing LapD2^C59S^ to export TelD into culture supernatants. Consistent with not playing an important role in LapD2 function, we found that an *S. intermedius* GC1825 *ΔlapD2* strain expressing plasmid-borne LapD2^C59S^ secretes wild-type levels of TelD (Fig 4F). Furthermore, the ability of LapD2 to form a heteromeric complex with LapD1 and TelD_LXG_ was unaffected by this mutation (Fig 4G). Finally, we noted that although LapD2 readily forms a Cys59 mediated cross-link when purified in isolation, this dimeric species is much less abundant when it is purified in complex with LapD1 and TelD_LXG_ (Fig 4E and 4G). Together, these data are indicative of the function of Lap2 proteins being less reliant on specific amino acids and more reliant on global aspects of protein structure.

### AlphaFold2 predicted models of Lap1 proteins reveals the location of a conserved FxxxD motif required for LXG effector secretion

Despite extensive efforts, we were unable to solve a crystal structure of a Lap1 protein. In general, we found that Lap1 proteins do not express well and were therefore poor candidates for crystallization experiments. Therefore, to better understand Lap1 function we used the recently released AlphaFold2 network to generate models of LapC1 and LapD1 (Fig 5A and Fig S5) (38). In parallel, we also ran AlphaFold2 on LapD2 and aligned the resulting model with our experimental crystal structure. As might be expected for a small single domain protein, the experimental and predicted models generally aligned well with a Cα RMSD of 2.0Å (Fig S5). However, we did note that the position of the turn region that connects the two α-helices occurs approximately seven residues earlier in the AlphaFold2 model compared to our crystal structure. Nonetheless, this result gave us reasonable confidence in the ability of AlphaFold2 to accurately predict the overall structure of Lap1 as members of this protein family share a similar size and predicted α-helical content as Lap2 proteins (Fig S1).

**Figure 5.**
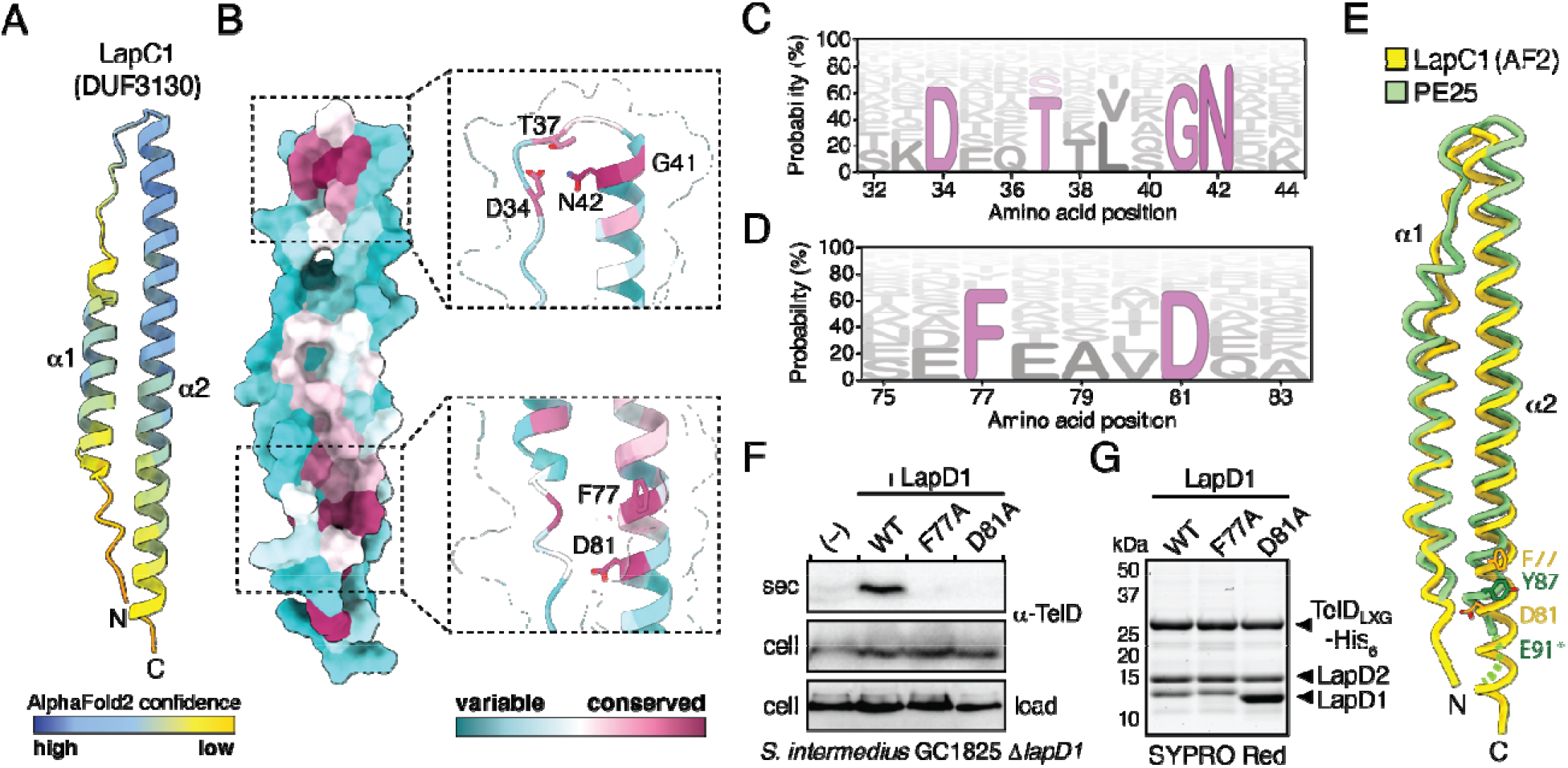
Lap1 modelling predicts a small α-helical protein harbouring a T7SSa export motif. (A) AlphaFold2 predicted structure of LapC1. Model is shown as a ribbon representation and coloured according to AlphaFold2 confidence level. (B) Surface representation of Lap1 sequence conservation mapped onto the LapC1 predicted structure. Sequences used for conservation analysis are available in Supplemental Table S2B. (C and D) HMM logo representation of the DxxTxxxGN and FxxxD sequence motifs identified in Lap1 family members. Probability is determined as a percent likelihood based on the Lap1 protein sequences in Supplemental Table S2B. (E and F) Mutation of the FxxxD motif in LapD1 blocks TelD secretion but does not impact TelD_LXG_-LapD1-LapD2 complex formation. Western blot analysis of the secreted and cell fractions of the indicated *S. intermedius* GC1825 strains (E). SYPRO Red staining of purified TelD_LXG_-LapD1-LapD2 wild-type and indicated variant complexes (F). (G) The FxxxD motif of Lap1 proteins is predicted to exist in a similar three-dimensional position as the YxxxE secretion signal of the T7SSa effector PE25. Noodle representation of LapC1 and PE25 (PDB ID: 4W4L) superposition. Structural models were aligned using the default matchmaker algorithm in ChimeraX. Asterisk indicates the approximate position of E91 as it was not modelled in the PE25 structure.

Overall, the AlphaFold2 generated Lap1 models adopt a helix-turn-helix arrangement similar to Lap2, with the exception of the first α-helix, which is markedly shorter than the second α-helix (Fig 5A and Fig S5). An alignment of 203 unique homologous Lap1 proteins was generated for LapC1 using one iteration of JackHMMER (Supplemental Table S2B and S4)(39). In contrast to Lap2, conservation mapping of Lap1 onto the predicted structure of LapC1 revealed two highly conserved regions within this protein family (Fig 5B). The first lies in the interhelical turn region and consists of a DxxTxxxGN motif (Fig 5B and 5C). We speculate that this motif is likely important for protein folding as the conserved residues face inwards towards one another and the side chains of Thr36 and Asn42 are predicted to hydrogen bond to one another based on their 2.9Å proximity. The second conserved region is solvent exposed, exists near the end of the second α-helix, and is punctuated by an FxxxD motif (Fig 5B and 5D). This motif drew our attention because it is remarkably similar to the YxxxD/E ‘export arm’ that serves as a secretion signal for Mycobacterial T7SSa effectors. Structural alignment of the predicted LapC1 structure with the crystal structures of the characterized Mycobacterial T7SSa effectors EspB and PE25 shows a striking overlap in the three-dimensional position of these residues, despite LapC1 possessing less than 15% sequence identity with either protein (Fig 5E and Fig S6).

Given our data suggesting that LapC1 itself is not secreted, we hypothesized that the FxxxD motif may act as an effector recognition signal that guides LXG proteins to the T7SSb when they are part of a Lap1-Lap2-LXG effector complex. To test this, the residues comprising this motif in LapD1 were targeted for site-specific mutagenesis to probe their role in TelD export. In line with functioning as a T7SSb export motif, we found that the secretion of TelD is not restored by plasmid-borne expression of LapD1^F77A^ or LapD1^D81A^ variants in an *S. intermedius* GC1825 *ΔlapD1* background (Fig 5F). To ensure that the observed lack of TelD secretion in these strains was not due to these site-specific variants compromising TelD-LapD1-LapD2 complex formation, we also introduced these mutations into our *E. coli* co-expression system and purified the LapD1 variant containing protein complexes. The results from this experiment demonstrate that LapD1^F77A^ or LapD1^D81A^ copurify with TelD_LXG_ and LapD2 in a manner that is comparable to wild-type LapD1 (Fig 5G). Collectively, these data show that the FxxxD motif of Lap1 proteins is not required for the formation of an effector pre-secretion complex but that it plays an essential role in LXG effector secretion by the T7SSb apparatus (Fig 6).

**Figure 6.**
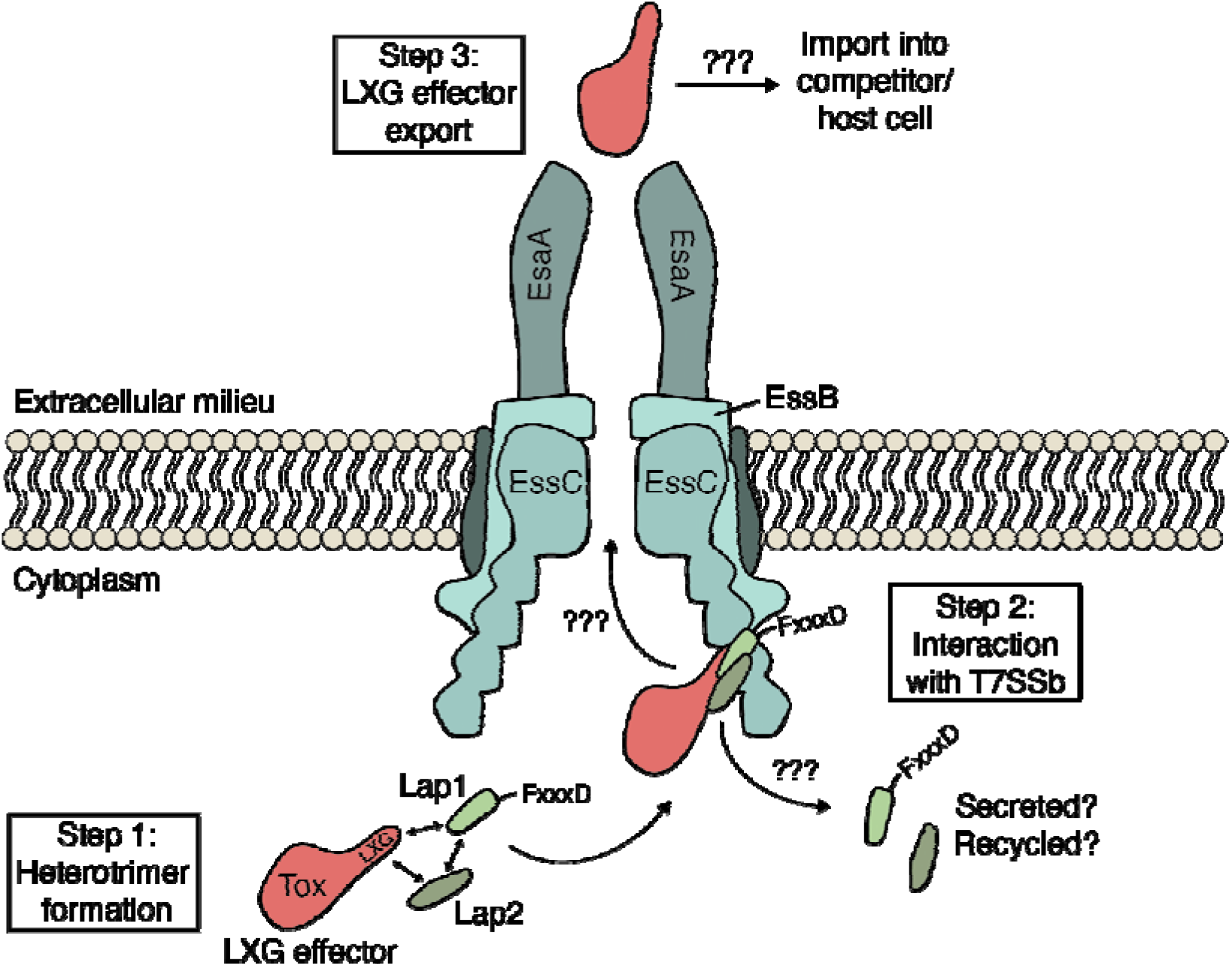
Model depicting LXG effector recruitment to the T7SSb apparatus by Lap1 and Lap2 targeting factors. Based on the findings described in this work, we propose that LXG effectors form a pre-secretion complex with cognate Lap1/DUF3130 and Lap2/DUF3958 proteins (step 1). The quaternary structure of this complex, in conjunction with the FxxxD motif found in Lap1 proteins, likely acts as a signal sequence that recruits LXG effectors to the T7SSb apparatus (step 2). The details of how T7SSb apparatuses facilitate protein export across the plasma membrane remain unknown but based on the findings of Rosenberg *et al*. on the ESX-1 T7SSa, this may involve effector-induced multimerization of EssC (step 3)(28). Once LXG effectors are released from the bacterial cell, those with cytotoxic activity enter the cytoplasm of their target cell by an unknown molecular mechanism.

## DISCUSSION

We have found that representative members of the Lap1/DUF3130 and Lap2/DUF3958 families of proteins function as targeting factors that promote the T7SSb-dependent secretion of cognate LXG effector proteins. Our structural and functional investigation also led us to discover that the former of these protein families possesses a critical sequence motif required for effector export. Altogether, these findings reveal several interesting parallels between the LXG-Lap1-Lap2 complexes defined herein and several well-characterized T7SSa effector families. For example, PPE proteins of *M. tuberculosis* are characterized by N-terminal domains of approximately the same size (~180-200 amino acids) and α-helical content as is predicted for LXG domains (21, 40). Moreover, these proteins are often encoded and expressed alongside members of the PE family of proteins, which like Lap1/Lap2, are ~100 amino acids in length, adopt a helix-turn-helix fold, and physically interact with their adjacently encoded effector (41). Several solved co-crystal structures of PE-PPE heterodimers demonstrates that these protein complexes form elongated α-helical bundles (42–45). Like Lap1 and Lap2, LXG domains are predicted to adopt an elongated α-helical structures and thus we speculate that LXG-Lap1-Lap2 heteromers may similarly adopt a side-by-side α-helical packing arrangement (21).

Another notable similarity between the PE and Lap1 protein families is the position of a conserved C-terminal motif, which in PE proteins and other T7SSa effectors is defined as Yxxx[D/E] whereas we identified a FxxxD motif in Lap1 proteins (29). In T7SSa effectors and EsxA proteins, this motif constitutes the so-called “export arm” and along with a WXG motif on a partner protein, functions as a bipartite secretion signal involved in the recruitment of effectors to the T7SS translocase EccC/EssC (46, 47). However, characterized PE proteins with this export motif are also typically co-secreted along with their partner PPE effector whereas we were unable to detect Lap1 or Lap2 in our secretion assays. While this finding could be due to the sensitivity of our measurements, it is also suggestive of a model in which these targeting factors dissociate from their cognate LXG effector during the secretion process. It is also interesting to note that some PE-PPE effector pairs also require a member of the globular EspG chaperone family to guide them to the T7SSa apparatus (48). While a globular chaperone, EsaE, has been shown to play a critical role in the T7SSb-dependent secretion of the non-LXG effector EsaD from *S. aureus*, a gene encoding a homologous protein does not exist in the *telC* and *telD* gene clusters (17).

The necessity of cognate Lap1-Lap2 targeting factors for the secretion of TelC and TelD is also interesting in the context of other LXG effectors. The TelC-producing B196 strain of *S. intermedius* secretes two additional LXG effectors named TelA and TelB, neither of which are encoded in gene clusters containing *lap1* or *lap2* homologous genes. TelA and TelB are instead encoded downstream of members of the DUF5082 and DUF5344 families of proteins, both of which are predicted helix-turn-helix proteins (34). Bacterial two-hybrid studies on the TelA- and TelB-associated DUF5082 proteins have shown that like Lap1 and Lap2, they physically interact with the LXG domain of their adjacently encoded LXG effector (18). Therefore, we speculate that these proteins likely play a similar role to the Lap1-Lap2 pairs described herein. The *S. aureus* effector TspA presents yet another intriguing case. In contrast to the LXG effectors of *S. intermedius*, the *tspA* operon consists of the effector gene followed by multiple copies of the immunity factor *tsaI* but no members of the small α-helical DUF families described above (21). This may indicate that the secretion of TspA does not require targeting factors for secretion or that TspA secretion requires the presence of small α-helical proteins encoded by ORFs found outside of the *tspA* gene cluster. Interestingly, gene clusters in *S. aureus* strains that encode the T7SSb apparatus often possess multiple genes encoding predicted small α-helical proteins, including EsxA, EsxB, EsxC and EsxD. All four of these proteins are secreted and either homo- or heterodimerize (49–51). Based on our findings, it is conceivable that one or more of these Esx proteins may function as targeting factors for TspA and/or other T7SSb effectors secreted by this bacterium.

While our identification and characterization of the factors required for LXG effector export has yielded new insight into the process of protein secretion by the T7SSb, future work structurally characterizing the identified three-protein complexes is required to better understand how LXG effector recognition by the T7SSb apparatus occurs at the molecular level. Studies on effector recognition by T7SSa pathways suggests that the EccC/EssC translocase may facilitate this recognition (28). However, more recent work on the T7SSb of *B. subtilis* found that LXG effectors directly interact with YukC (EssB), a protein that the authors of this study propose serves as the central interaction hub that holds the T7SSb apparatus together (24). Regardless of which apparatus protein(s) recognise LXG effectors, our data suggests that the ‘signal sequence’ that allows for this recognition is likely defined by the quaternary structure of LXG-Lap1-Lap2 complexes and the FxxxD export motif found within Lap1. Upon interaction with the apparatus, we hypothesize that LXG effector export is facilitated by a conformational change in the T7SSb structure that is energetically linked to ATP binding and hydrolysis by EssC. Effectors are then transported through the cell envelope in a single step via a protein channel comprised of the various T7SSb structural subunits, the molecular details of which remain obscure. Several recent cryo-EM structures of Mycobacterial T7SSa apparatuses have provided profound mechanistic insights into the function of T7SSa pathways and it is probable that structures of the T7SSb will similarly inform our understanding of protein export by this complex molecular machine (52–55).

## MATERIALS AND METHODS

### Bacterial strains, plasmids, and growth conditions

*S. intermedius* strains used in this study were generated from either the B196 or GC1825 wild-type strains and genomic DNA isolated from these strains was used for molecular cloning. *E. coli* XL1-blue was used for molecular cloning and plasmid maintenance. *E. coli* BL21 (DE3) CodonPlus and B834 (DE3) were used for protein expression of native and selenomethionine substituted proteins, respectively. The complete list of bacterial strains generated for this study can be found in Supplemental Table S3. *E. coli* overexpression was performed using the IPTG-inducible pETDuet-1 and pET29b vectors, while pDL277 was used for constitutive gene expression in *S. intermedius*. PCR amplification of genes of interest for this study was done with Phusion polymerase (NEB). For pET vector cloning the PCR amplicons were digested with restriction endonucleases NdeI/XhoI for pET29b/pETduet-1 MCS2 or BamHI/SalI for pETduet-1 MCS1. DNA ligation was then done using T4 DNA ligase. These constructs were cloned with N- or C-terminal His-6 tags to facilitate affinity purification as required. Cloning into the pDL277 vector was done with restriction endonucleases BamHI/SalI followed by ligation with T4 DNA ligase. In this case, *S. intermedius* genes were fused with the P96 promoter of *Streptococcus pneumoniae* by <u>s</u>plicing by <u>o</u>verlap <u>e</u>xtension (SOE) PCR as previously described (18). A complete list of the plasmids used in this study can be found in Supplemental Table S4. *E. coli* was grown in lysogeny broth at 37°C at 225rpm. 50ug/mL kanamycin and 150ug/mL carbenicillin was added to the media when growing strains with the pET29b and pETduet-1 vectors, respectively. *S. intermedius* strains were grown in Todd Hewitt broth supplemented with 0.5% yeast extract at 37°C and 5% CO_2_ without shaking. 50ug/mL of spectinomycin for *S. intermedius* or 100ug/mL of spectinomycin for *E. coli* was added to media when growing strains with the pDL277 plasmid. For all *S. intermedius* experiments, strains were first grown on solid media before being inoculated into liquid culture to ensure consistent growth between strains.

### DNA manipulation

*S. intermedius* B196 and GC1825 genomic DNA was prepared by using InstaGene Matrix (Bio-Rad) to extract and purify DNA from 2 mL of cells pelleted from an overnight culture. Primers used in this study were synthesized by Integrated DNA Technology (IDT). Molecular cloning was performed using Phusion polymerase, appropriate restriction enzymes, and T4 DNA ligase (NEB). All Sanger sequencing was performed by Azenta.

### Transformation of *S. intermedius*

*S. intermedius* B196 and GC1825 strains were back diluted 1:10 from an overnight culture, grown to OD_600_ = 0.5, and supplemented with 5uL of 0.1mg/mL competence stimulating peptide (DSRIRMGFDFSKLFGK, synthesized by Genscript). Cultures were then incubated at 37°C and 5% CO_2_ without shaking for 45 minutes (GC1825) or two hours (B196). Approximately 100ng of plasmid or linear DNA was then added and the cultures were again incubated for three hours (1 hour for GC1825). 100uL of these cultures were then plated on Todd Hewitt plates supplemented with 0.5% yeast extract and either 50 ug/mL spectinomycin to select for pDL277 transformants or 250 ug/mL kanamycin for allelic replacement mutants.

### Gene deletion in *S. intermedius* by allelic replacement

Our *S. intermedius* gene deletion protocol was previously described in (56). In brief, deletion constructs were made using SOE PCR to fuse a spectinomycin promoter to a kanamycin resistance cassette flanked by two 1000bp fragments of DNA that are immediately adjacent to the target gene. These constructs were cloned into pETduet-1 with the final plasmid designation being pETduet-1::5’geneflank_SpecProm_*kanR*_3’geneflank. Plasmids were then digested with BamHI and NotI and the deletion fragment was gel extracted (Monarch DNA gel extraction kit, NEB). 100ng of purified deletion fragment was then added to competent *S. intermedius* cells and mutants were selected for by plating on Todd Hewitt agar with 0.5% yeast extract and 250 ug/mL kanamycin. All gene deletions were confirmed by colony PCR.

### Secretion assays

20mL cultures of *S. intermedius* were grown overnight to an OD_600_=1.0. Cell and supernatant fractions were then separated by centrifugation at 4000g for 15 minutes and cell fractions were washed once in PBS pH 7.4 before being resuspended in 100 uL of PBS. 100 uL of Laemmli buffer was added and samples were boiled for 10 minutes. Supernatant fractions were incubated at 4°C overnight after adding trichloroacetic acid to a final concentration of 10%. Precipitated proteins were then centrifuged at 35,000g for 30 minutes and the resulting pellets were washed once with cold acetone. The pellets were then centrifuged at 35,000g for an additional 30 minutes and the acetone was decanted off. Any remaining acetone was left to evaporate off in a fume hood. The dry pellets were then resuspended in minimal Laemmli buffer diluted with urea (300uL 4X Laemmli, 600uL 8M urea) and boiled for 10 minutes. Both the cell and secreted samples were analysed using SDS PAGE gels run with a tris-tricine based running buffer (see below) and Western blot analysis.

### Antibody generation

A custom polyclonal antibody for the TelD protein was generated for this study. In brief, the LXG domain of TelD (amino acids 1-203) with a C-terminal His_6_ tag was co-expressed with LapD1 and purified by affinity and size exclusion chromatography as described (see “protein expression and purification”) except with PBS pH 7.4 in place of Tris-HCl pH 8.0 as the buffer system. In total, 10mg of purified protein was shipped to Genscript for custom polyclonal antibody production. Western blots on samples prepared from the appropriate *S. intermedius* GC1825 deletion strains determined that this antibody specifically recognizes TelD but not LapD1. Generation of the α-TelC and α-EsxA antibodies have been described previously (18, 56).

### SDS-PAGE, SYPRO red staining and Western blotting

SDS-PAGE gels run for this study were done using a tris-tricine buffer system (200mM Tris, 100mM Tricine, 0.1% SDS, pH 8.3) to better resolve low molecular weight proteins (<20kDa)(57). Protein visualization on SDS-PAGE gels was done with the SYPRO Red protein gel stain (Invitrogen). The gel was rinsed briefly in DI water before being stained for one hour with 1:5000 SYPRO Red (Invitrogen) diluted in 10% (v/v) acetic acid. The gel was then destained for 15 minutes in 7.5% (v/v) acetic acid before being imaged on a Chemidoc imaging system (Bio-Rad). For western blots, the resolved proteins were transferred to a nitrocellulose membrane by wet transfer (100V, 30 minutes). Nitrocellulose membranes were then blocked with 5% skim milk dissolved in TBS-T for 30 minutes with light agitation followed by addition of primary antibody (titer 1:5000) to the blocking buffer and further incubation for 1 hour. Blots were washed for five minutes three times with TBS-T then incubated in TBS-T with an HRP-conjugated anti-rabbit secondary antibody (titer 1:5000) for 45 minutes. After three additional five-minute washes, the blots were developed using Clarity Max Western ECL reagent (Bio-Rad) and imaged with a ChemiDoc XRS+ (Bio-Rad).

### Co-immunoprecipitation in *Streptococcus intermedius*

Co-immunoprecipitation assays were performed on VSV-G tagged TelC in a *ΔtelC-tipC2* background (ΔSIR_1486-1489) and VSV-G tagged LapC1 in a *ΔlapC1* background (ΔSIR_1491). In both experiments, strains lacking SIR1486-1489 or SIR1491 but containing empty pDL277 were used as negative controls. 50mL cultures of *S. intermedius* were grown to an OD of 0.5 and centrifuged at 5000g for 15 minutes to harvest cells. The pellets were then resuspended and incubated in lysis buffer (20mM Tris-HCl pH 7, 150mM NaCl, 10% glycerol, 5 mg/mL lysozyme, 100U/mL mutanolysin, 1 mM PMSF) and incubated at 37°C for 30 minutes. Cells were lysed by sonication (three, thirty second pulses at 30 amps) and the cell pellets were removed by centrifugation at 30,000g for 30 minutes at 4°C. The supernatants were then transferred to fresh 2 mL Eppendorf tubes and incubated with 50uL of anti-VSV-G beads overnight at 4°C with gentle agitation. The beads were harvested by centrifugation at low speed (<100g) and washed thrice with 10 mLs of wash buffer (20mM Tris-HCl pH 7, 150mM NaCl, 10% glycerol). An additional three wash steps were performed with 50 mM ammonium bicarbonate. The beads were then covered in a minimal amount of ammonium bicarbonate buffer and the bound protein was digested with 10 ng/ul of sequencing grade trypsin for four hours at 37°C. The buffer was then harvested, and the beads were washed with an additional 50uL of ammonium bicarbonate buffer to remove any remaining peptides. The peptide samples were then incubated with 1 mM tris(2-carboxyethyl)phosphine for one hour at 37°C to reduce any disulphide bonds. Iodoacetamide was added to a final concentration of 10 mM and the samples were incubated in the dark at room temperature for 30 minutes. This reaction was quenched with 12 mM N-acetylcysteine. The peptides were purified using Pierce C18 spin columns (Thermo Scientific). LC-MS/MS analysis of the purified peptides was done at the Sick Kids Proteomics, Analytics, Robotics, and Chemical Biology Centre (SPARC) at The Hospital for Sick Children.

### TelD toxicity assay

*E. coli* XL1 blue was transformed with either the pSCRhaB2 plasmid encoding the *telD* toxin gene or an empty vector control. For the toxicity plating assay, these strains were OD matched and serially diluted (1:10) then plated on LB plates containing 200 μg/mL of trimethoprim with and without 0.1% L-rhamnose. The plates were incubated at 37°C overnight and then imaged using an iPhone 11 (Apple). Growth curves were generated by back diluting overnight cultures 1:100 into fresh LB media supplemented with 200 μg/mL trimethoprim and 15 μg/mL gentamicin in a 96-well plate. The cell cultures were allowed to grow at 37°C with shaking for 1.5 hours at which point toxin expression was induced by adding L-rhamnose to a final concentration of 0.1% and immunity protein expression was induced by adding IPTG to a final concentration of 0.1 mM. The OD of the cultures was measured with a Synergy 4 Microplate Reader (Biotek Instruments).

### Protein expression and purification

All native proteins were expressed in *E. coli* BL21(DE3) CodonPlus whereas selenomethionine-labeled LapD2 was expressed in *E. coli* B834 (DE3). In general, protein expression strains were grown in LB in a shaking incubator at 37°C to an OD_600_=0.5. Temperature was then lowered to 18°C and protein expression was induced with 1mM IPTG followed by overnight protein expression (approximately 18 hours). Cells were then centrifuged and lysed by sonication (four pulses, 30% amplitude, 30 seconds) in lysis buffer (20mM Tris-HCl pH 8.0, 300mM NaCl, 10mM imidazole). Cellular debris was cleared from the lysate by centrifugation at 35,000g for 30 minutes and the lysate was run over Ni-NTA resin using a gravity flow column on the benchtop. Resin was then washed three times with 20mL lysis/wash buffer and protein was eluted in 4mL of elution buffer (20mM Tris-HCl pH, 8.0, 300mM NaCl, 400mM imidazole). Eluted protein was further purified by size exclusion chromatography using a HiLoad 16/600 Superdex 200 connected to an ÄKTAexplorer (Cytiva). Selenomethionine-labeled protein was similarly expressed using *E. coli* B834 (DE3) except that the cells were grown in SelenoMethionine Media (Molecular Dimensions) supplemented with 40 mg/L of L-selenomethionine.

### Protein Crystallization

Native and selenomethionine-labeled LapD2 was concentrated to 10 mg/ml and screened for crystallization conditions using the MCSG1-4 crystallization suites (Anatrace) and the hanging drop vapour diffusion method. After one week, trapezoid shaped crystals formed in 0.2M lithium sulfate, 0.1M Tris-HCl, pH 8.0, 30% (w/v) PEG 4000. Crystals were cryoprotected using a buffer identical to the crystallization buffer but supplemented with 20% ethylene glycol.

### X-ray data collection, structure determination and model refinement

X-ray data were collected with the Structure Biology Center sector 19-ID at the Advanced Photon Source. Diffraction of both selenomethionine-incorporated and native protein crystals were measured at a temperature of 100 K using a 0.3s exposure and 0.5 degree of rotation over 450°. Native and selenomethionine-incorporated crystals diffracted to resolutions of 2.20 Å and 2.42 Å, respectively, and the diffraction images were collected on a dectris Pilatus 3 X 6M detector with an X-ray wavelength near the selenium edge of 12.66 keV (0.97926 Å). Diffraction data were processed using the HKL3000 suite (58). The structure of LapD2 was determined by SAD phasing with data from selenomethionine-containing protein crystal using SHELX C/D/E (59), mlphare and dm (60), and initial automatic protein model building with Buccaneer (61), all implemented in the HKL3000 software package (58). The initial model of the structure of the homodimer was completed manually by using Coot (62) and briefly refined using refmac (63). Using this dimeric structure from the SAD phasing as the search model, molecular replacement was applied with the native data using molrep implemented in HKL3000. The structure was then refined iteratively using Coot for manual adjustment and Phenix (phenix.refine)(64) for restrained refinement until *R*_work_ and *R*_free_ values converged to 0.23 and 0.26, respectively. The final refined structure contained two copies of homodimeric LapD1 with each dimer formed through a disulfide bond. The stereochemistry of the structure was assessed using PROCHECK (65) and a Ramachandran plot and was validated using the PDB validation server. X-ray data and refinement statistics are listed in Table 1. All structural figures were generated using UCSF ChimeraX (66).

### Protein structure prediction and analysis

Surface hydrophobicity (67), conservation mapping (68), structural alignments were visualized using ChimeraX’s built in functions with default parameters (69). DALI pairwise alignment was used to calculate reported RMSD values (37). 2D protein structure predictions were generated by PSIPRED 4.0 on the UCL PSIPRED Workbench (70) (http://bioinf.cs.ucl.ac.uk/psipred/). 3D Protein structure predictions were performed by AlphaFold v2.0.0 running on our local server with default parameters (38).

### Sequence analysis, conservation mapping and sequence logos

Homologous sequences to LapC1, LapC2, LapD1 and LapD2 were identified using JackHMMER (HmmerWeb version 2.41.2) searches of the UniprotKB database, restricted to the phylum Firmicutes, iterating until at least 100 sequences were obtained (39). Accessions were downloaded and full sequences of active entries were subsequently retrieved from Uniprot. Duplicate sequences were removed and the remaining aligned using MAFFT (scoring matrix: BLOSUM30)(71) implemented in Geneious Prime 2022.1.0 (www.geneious.com). Final sequence lists used for HMM logo generation can be found in Supplemental Tables S2A, S2B, and S2C. HMMs were generated and initially visualized by uploading multiple sequence alignments to the Skylign webserver (www.skylign.org) and set to “create HMM – remove mostly empty columns” (72). The resulting matrices were downloaded as tabular text, formatted and then visualized using Logomaker (73). Sequence alignments depicted in Supplemental Figures 3, 4 and 6 were generated using M-Coffee on the T-Coffee webserver (https://tcoffee.crg.eu) (74) and visualized with the ESPript 3.0 webserver (75) (https://espript.ibcp.fr/ESPript/ESPript/).

## DATA AVAILABILITY

The atomic coordinates and structure factors for LapD2 have been deposited in the Protein Data Bank under accession code 7UH4.

Scripts and intermediate files used to generate figures are available at: https://github.com/dirkgreb/Lap_paper_2022

## SUPPORTING INFORMATION

This article contains supporting information.

## ACKNOWLEDGEMENTS

The authors would like to thank Andrew McArthur and Amos Raphenya for setting up AlphaFold2 on their Apollo server and Shehryar Ahmad, Nathan Bullen, and Andrea Alexei for their constructive feedback on the manuscript. We also thank the SickKids Proteomics, Analytics, Robotics & Chemical Biology Centre (SPARC) at the Hospital for Sick Children in Toronto, Ontario for mass spectrometry experiments, and the members of the Structural Biology Center (SBC) at Argonne National Laboratory for their help with data collection at the 19-ID beamline. The use of the SBC beamlines at the Advanced Photon Source is supported by the U.S. Department of Energy (DOE) Office of Science and operated for the DOE Office of Science by Argonne National Laboratory under contract no. DE-AC02-06CH11357.

## AUTHOR CONTRIBUTIONS

T.A.K. and J.C.W conceived the study. All authors contributed to experimental design. M.G.S. provided the *S. intermedius* GC1825 strain and performed whole genome sequencing on this strain. T.A.K., O.M., and J.C.W. generated strains and plasmids. T.A.K. and P.Y.S. expressed, purified, and crystallized protein. T.A.K., D.W.G., Y.K., and J.C.W. solved and analyzed the crystal structure. T.A.K. and O.M. performed biochemical experiments. T.A.K., D.W.G., and J.C.W. analyzed the data. T.A.K., D.W.G. and J.C.W. wrote the paper. All authors provided feedback on the manuscript.

## FUNDING AND ADDITIONAL INFORMATION

The use of SBC beamlines at the Advanced Photon Source is supported by the U.S. Department of Energy (DOE) Office of Science and operated for the DOE Office of Science by Argonne National Laboratory under contract no. DE-AC02-06CH11357. T.A.K. is supported by an Alexander Graham Bell Canada Graduate Scholarship from the Natural Sciences and Engineering Research Council of Canada (NSERC). This work was supported by a project grant (PJT-173486) from the Canadian Institutes of Health Research (CIHR). M.G.S is the Canada Research Chair in Interdisciplinary Microbiome Research. J.C.W. is the Canada Research Chair in Molecular Microbiology and holds an Investigators in the Pathogenesis of Infectious Disease Award from the Burroughs Wellcome Fund.

## SUPPLEMENTAL FIGURE & TABLE LEGENDS

**Figure S1. Secondary structure predictions for EsxA, LapC1 and LapC2 from *Streptococcus intermedius* B196.** Graphical output from PSIPRED 4.0 analyses of EsxA, LapC1 and LapC2. Per-residue secondary structure predictions and confidence scores are indicated above each amino acid in the sequence.

**Figure S2. Sequence and predicted secondary structure alignment of TelD and TspA.** Secondary structure assignments are based on AlphaFold 2 predicted tertiary structures. Overall pairwise sequence identity is 23.8%. TelD and TspA have highest levels of sequence homology within their predicted N-terminal LXG domains. Dashed blue box indicates each effector’s LXG motif.

**Figure S3. Toe-to-Toe packing arrangement of LapD2 and structural alignment of LapD2 to *M. tuberculosis* EsxB.** (A-B) LapD2 chains A and C interact with one another in a toe-to-toe manner that involves both N- and C-termini. (C) Structural alignment of LapD2 with *M. tuberculosis* EsxB (PDB code 3FAV) shown in ribbon representation.

**Figure S4. Sequence alignment of LapD2, LapC2 and four additional Lap2 homologs, and sequence logo representation of Lap2 regions that possess sequence conservation.** (A) Multiple sequence alignment of LapD2 with four randomly selected homologs identified by JackHMMER (UniprotKB accessions listed), and LapC2. (B) Normalized HMM logos generated from the entire JackHMMER sequence hit table reveal a high degree of sequence variability across the group, even in the most conserved regions of the protein (3 and 4).

**Figure S5. AlphaFold2 predicted structure and sequence conservation mapping of LapD1 and comparison of the LapD2 crystal structure to its AlphaFold2 model.** (A) AlphaFold2 model of LapD1 coloured by confidence score. (B) Surface representation of Lap1 sequence conservation mapped onto the LapD1 predicted structure. (C) LapD2 crystal structure (gold) aligned to the AlphaFold2 predicted model (coloured by confidence score, as in panel A). (D) AlphaFold-multimer models of LapC1 and LapC2 in hypothetical homodimeric (left and right panels, respectively) and heterodimeric (middle) arrangements.

**Figure S6. AlphaFold2 predicted structure of LapC1 aligned to crystal structures of the Type VIIa substrates EspB and PE25.** (A) Predicted structure of LapC1 (yellow) aligned to the Y-subdomain (dark green) of EspB from *M. tuberculosis* (PDB ID: 4WJ1) reveals a conserved FxxxD motif found in a similar location as the YxxxD/E export motif required for EspB secretion. (B) Structural alignment of predicted LapC1 structure to PE25 when in complex with its cognate PPE41 protein and EspG5 chaperone (PDB ID: 4W4L). (C-D) Pairwise sequence alignments of LapC1 to EspB_1-96_ (C) and PE25 (D).

**Table S1. Spectral counts for TelC-V and LapC1-V immunoprecipitated samples and their respective control samples.** Mass-spectrometry of the TelC-V and LapC1-V immunoprecipitation experiments was performed at the SPARC facility at the Hospital for Sick Children. Accession numbers and IDs are given along with the common name for the top 100 proteins identified in our experiments. Proteins of interest are listed in order of total counts (between all four samples). The TelC experimental/control counts are displayed in light grey columns, while the LapC1 experimental/control counts are displayed in dark grey.

**Table S2. Accession codes and sequence information for LapD2, LapC1, and LapD1 homologs identified with three iterations of JackHMMER.** List of homologous proteins for (A) LapD2, (B) LapC1, and (C) LapD1 are listed with their entry codes, protein names, gene names, organism, and length (in amino acids). These lists were generated by three, one, and one iteration(s) of a JackHMMER homology search, respectively.

**Table S3. Strains used in this study.**

**Table S4. Plasmids used in this study.**

