## Supplemental Figures for "Dual targeting factors are required for LXG toxin export by the bacterial type VIIb secretion system"

**Figure S1. Secondary structure predictions for EsxA, LapC1 and LapC2 from *Streptococcus intermedius* B196.**

**Figure S2. Sequence and predicted secondary structure alignment of TelD and TspA.**

**Figure S3. Toe-to-Toe packing arrangement of LapD2 and structural alignment of LapD2 to *M. tuberculosis* EsxB.**

**Figure S4. Sequence alignment of LapD2, LapC2 and four additional Lap2 homologs, and sequence logo representation of Lap2 regions that possess sequence conservation.**

**Figure S5. Alphafold2 predicted structure and sequence conservation mapping of LapD1 and comparison of the LapD2 crystal structure to its Alphafold2 model.**

**Figure S6. Alphafold2 predicted structure of LapC1 aligned to crystal structures of the Type VIIa substrates EspB and PE25.**

**Supplemental references**

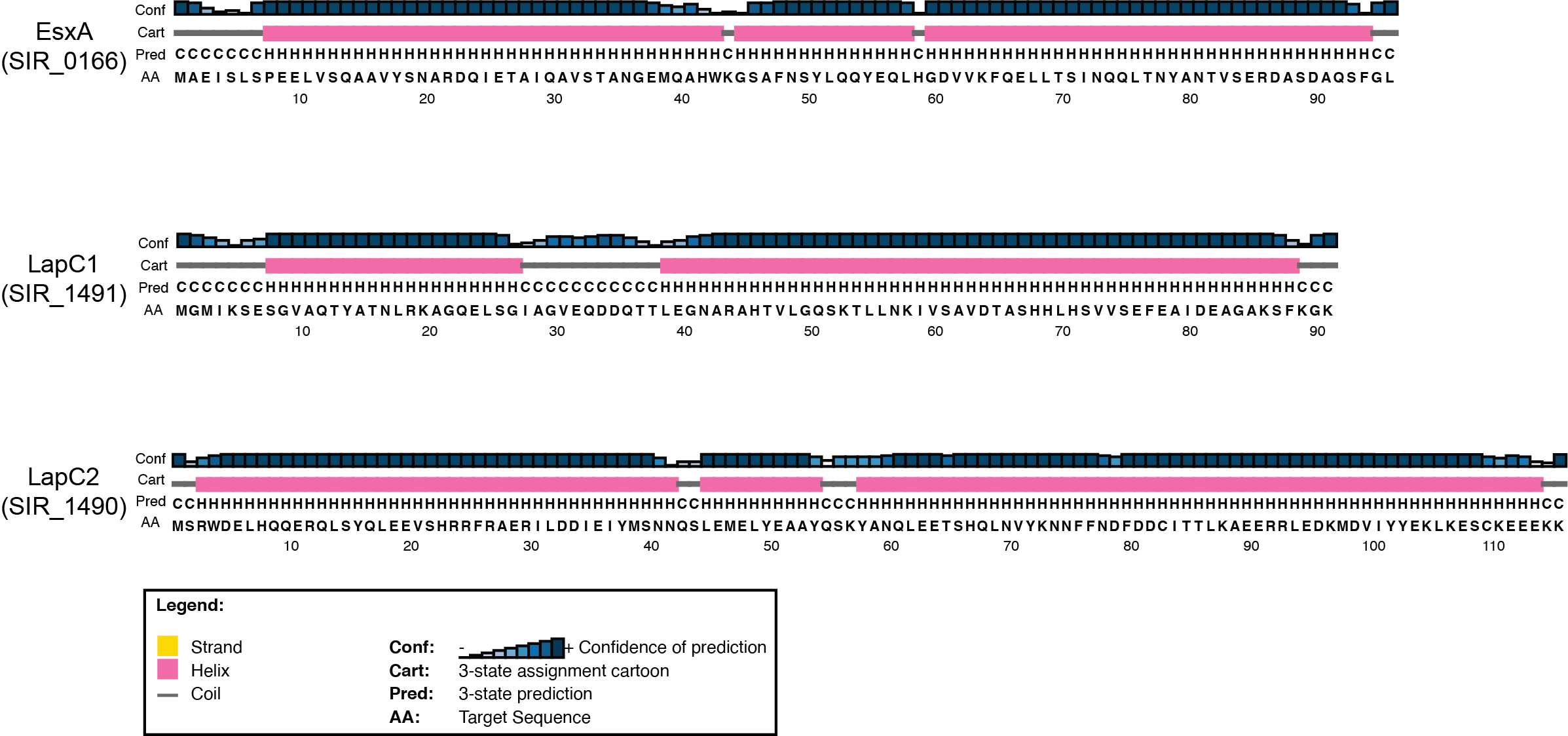

**Figure S1. Secondary structure predictions for EsxA, LapC1 and LapC2 from *Streptococcus intermedius* B196.** Graphical output from PSIPRED 4.0 analyses of EsxA, LapC1 and LapC2. Per-residue secondary structure predictions and confidence scores are indicated above each amino acid in the sequence.

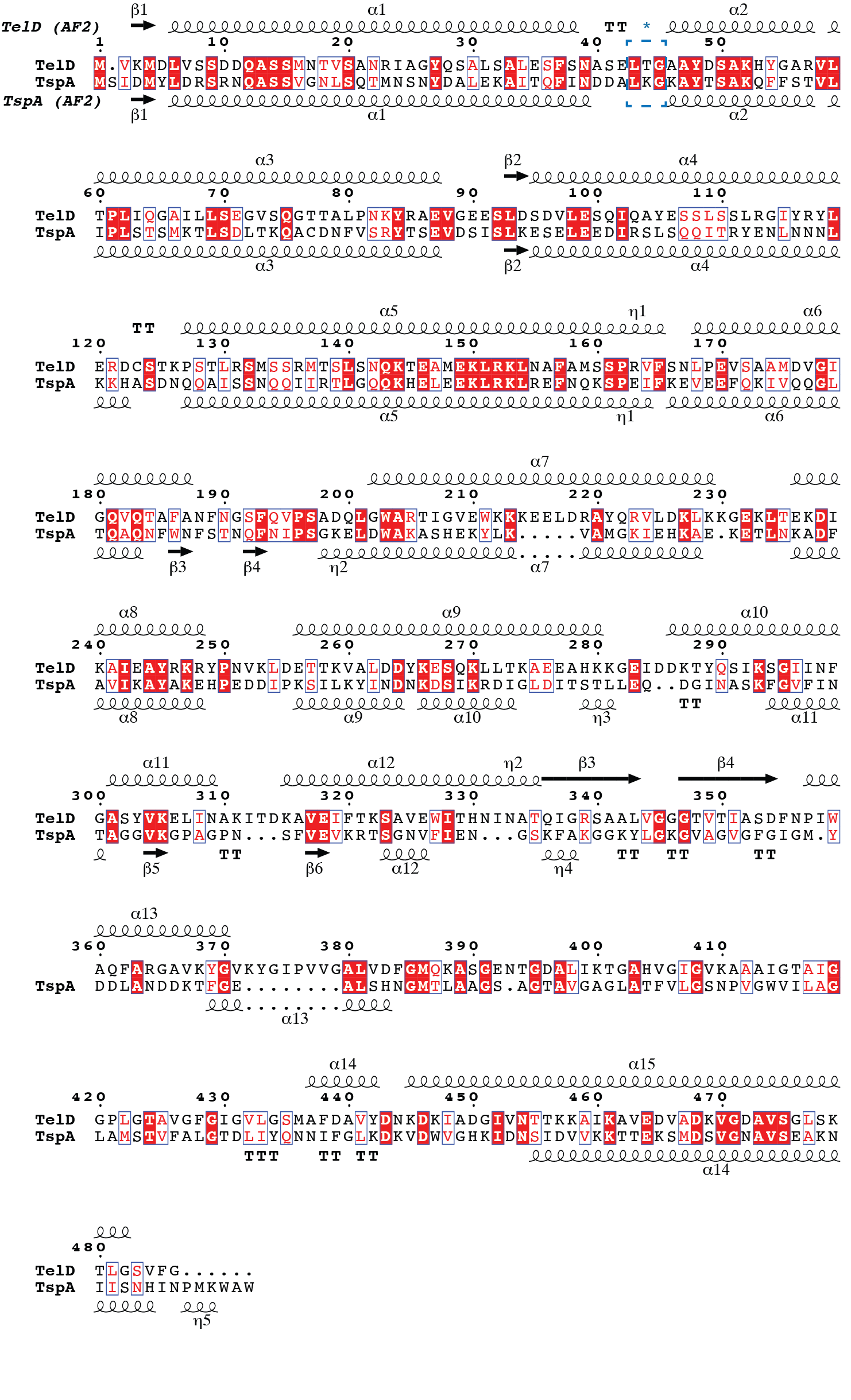

**Figure S2. Sequence and predicted secondary structure alignment of TelD and TspA.** Secondary structure assignments are based on Alphafold 2 predicted tertiary structures. Overall pairwise sequence identity is 23.8%. TelD and TspA have highest levels of sequence homology within their predicted N-terminal LXG domains. Dashed blue box indicates each effector’s LXG motif.

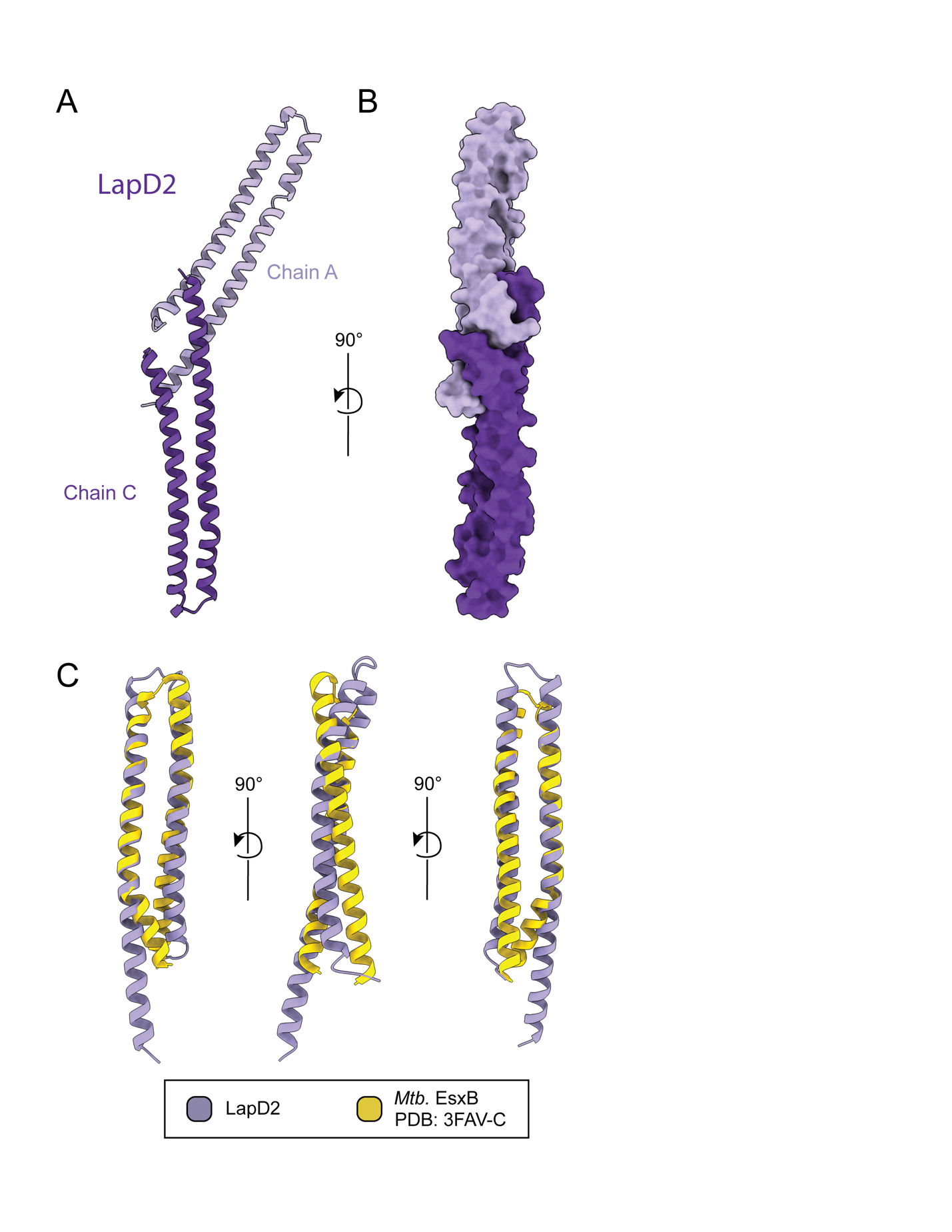

**Figure S3. Toe-to-Toe packing arrangement of LapD2 and structural alignment of LapD2 to *M. tuberculosis* EsxB.** (A-B) LapD2 chains A and C interact with one another in a toe-to-toe manner that involves both N- and C-termini. (C) Structural alignment of LapD2 with *M. tuberculosis* EsxB (PDB code 3FAV) shown in ribbon representation.

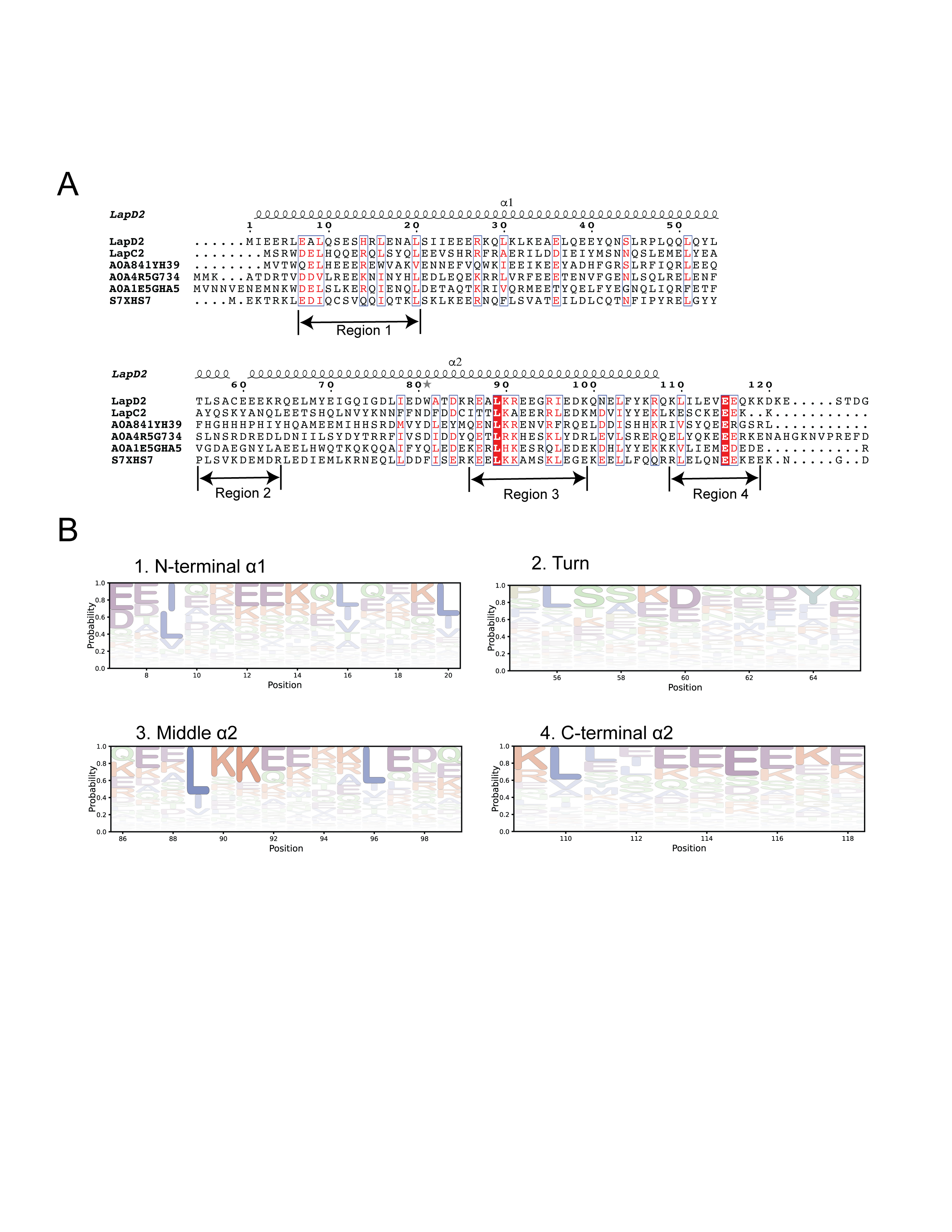

**Figure S4. Sequence alignment of LapD2, LapC2 and four additional Lap2 homologs, and sequence logo representation of Lap2 regions that possess sequence conservation.** (A) Multiple sequence alignment of LapD2 with four randomly selected homologs identified by JackHMMER (UniprotKB accessions listed), and LapC2. (B) Normalized HMM logos generated from the entire JackHMMER sequence hit table reveal a high degree of sequence variability across the group, even in the most conserved regions of the protein (3 and 4).

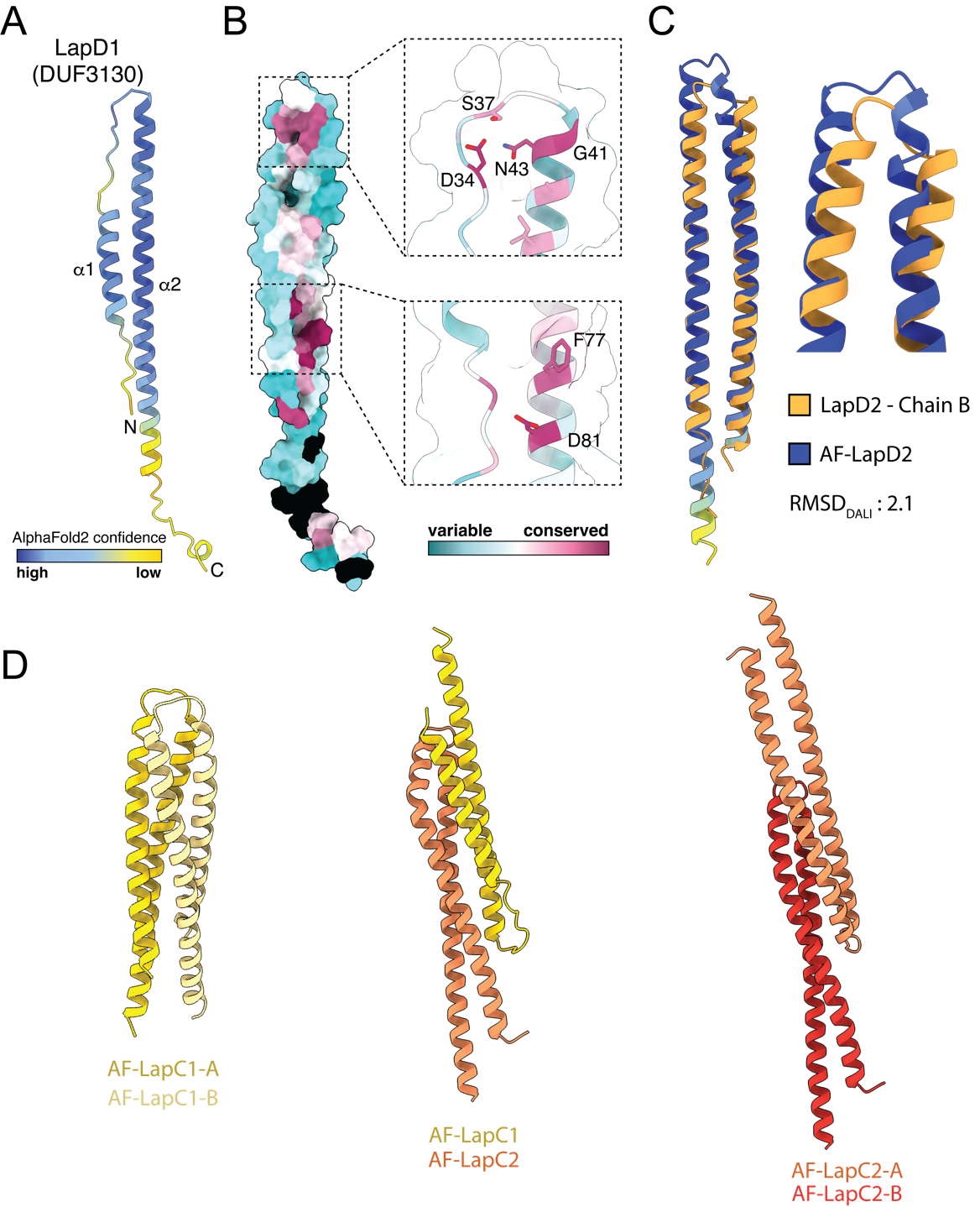

**Figure S5. AlphaFold2 predicted structure and sequence conservation mapping of LapD1 and comparison of the LapD2 crystal structure to its AlphaFold2 model.** (A) AlphaFold2 model of LapD1 coloured by confidence score. (B) Surface representation of Lap1 sequence conservation mapped onto the LapD1 predicted structure. (C) LapD2 crystal structure (gold) aligned to the AlphaFold2 predicted model (coloured by confidence score, as in panel A). (D) AlphaFold-multimer models of LapC1 and LapC2 in hypothetical homodimeric (left and right panels, respectively) and heterodimeric (middle) arrangements.

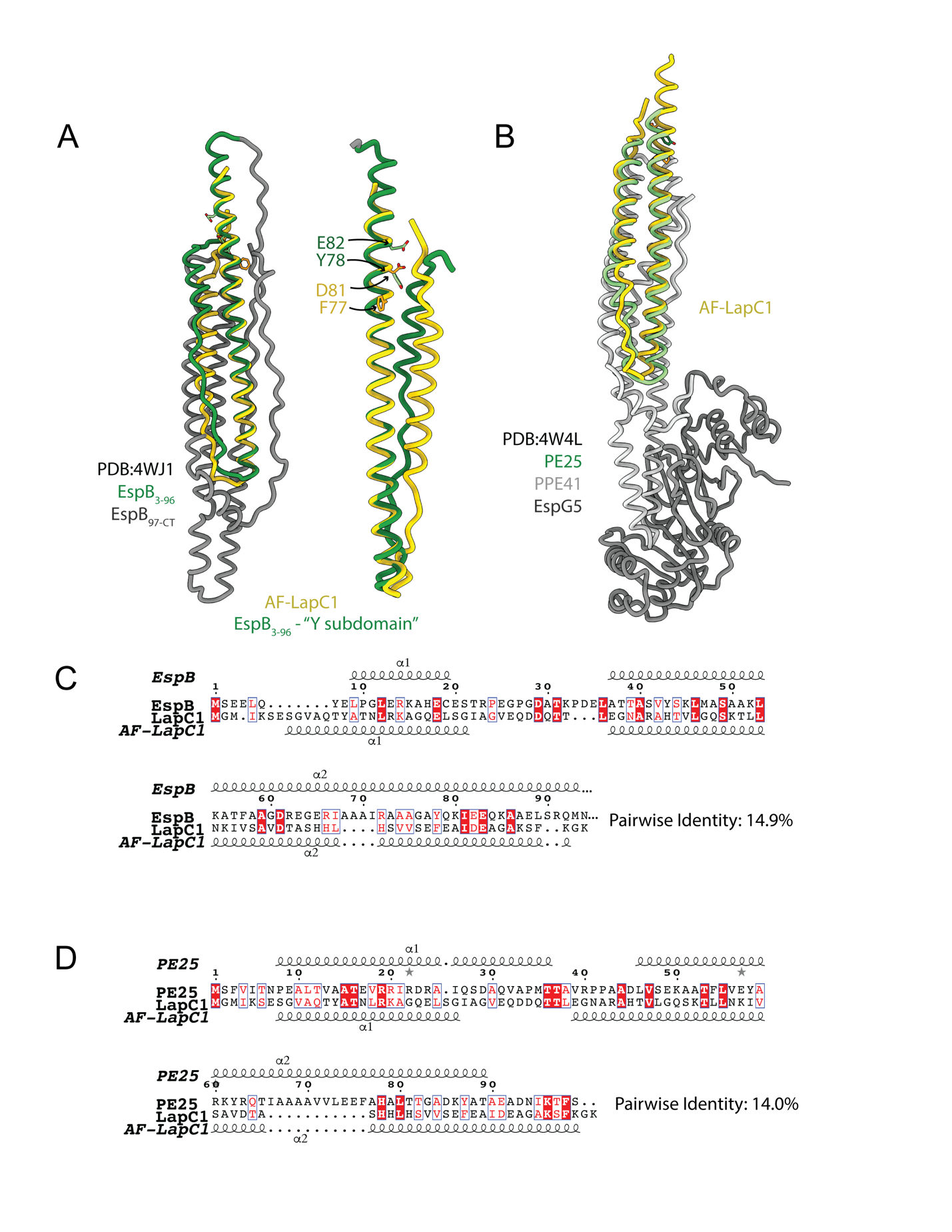

**Figure S6. Alphafold2 predicted structure of LapC1 aligned to crystal structures of the Type VIIa substrates EspB and PE25.** (A) Predicted structure of LapC1 (yellow) aligned to the Y-subdomain (dark green) of EspB from *M. tuberculosis* (PDB ID: 4WJ1) reveals a conserved FxxxD motif found in a similar location as the YxxxD/E export motif required for EspB secretion. (B) Structural alignment of predicted LapC1 structure to PE25 when in complex with its cognate PPE41 protein and EspG5 chaperone (PDB ID: 4W4L). (C-D) Pairwise sequence alignments of LapC1 to EspB_1-96_ (C) and PE25 (D).
