## Supplemental Table S1 for "Dual targeting factors are required for LXG toxin export by the bacterial type VIIb secretion system"

**Table S1. Spectral counts for TelC-V and LapC1-V immunoprecipitated samples and their respective control samples.**

| **#** | **Identified Proteins** | **Accession Number** | **Alternate ID** | **Δ*telC* ctrl** | **Δ*telC* + *telC*-V** | **Δ*wxgC* ctrl** | **Δ*wxgC*+*wxgC-*V** |
| --- | --- | --- | --- | --- | --- | --- | --- |
| 1 | TelC-VSV-G tagged | TelC_VSV-G |  | 0 | 716 | 11 | 191 |
| 2 | WxgC-VSV-G tagged | WxgC_VSV-G |  | 0 | 288 | 2 | 184 |
| 3 | Enolase | T1ZFF4 | eno | 35 | 45 | 28 | 47 |
| 4 | Polyribonucleotide nucleotidyltransferase | T1ZFD6 | pnpA | 39 | 37 | 35 | 36 |
| 5 | Uncharacterized protein | T1ZGI6 | SIR_1490 | 0 | 73 | 0 | 28 |
| 6 | Oligopeptide-binding protein AmiA | T1ZFZ2 | amiA | 21 | 25 | 12 | 15 |
| 7 | 60 kDa chaperonin | T1ZGB6 | groL | 20 | 20 | 8 | 15 |
| 8 | ABC-type transport system, periplasmic binding protein | T1ZEB9 | SIR_1223 | 13 | 16 | 9 | 11 |
| 9 | Uncharacterized protein | T1ZDT0 | SIR_1033 | 13 | 10 | 6 | 10 |
| 10 | Putative extracellular solute-binding protein | T1ZFL7 | SIR_1387 | 11 | 11 | 7 | 11 |
| 11 | Chaperone protein DnaJ | T1ZG02 | dnaJ | 10 | 9 | 12 | 13 |
| 12 | ABC transporter, substrate-binding protein | T1ZG31 | SIR_1454 | 9 | 13 | 5 | 10 |
| 13 | Isoprenyl transferase | T1ZGQ4 | uppS | 5 | 13 | 7 | 8 |
| 14 | Uncharacterized protein | T1ZGR7 | SIR_1322 | 7 | 13 | 2 | 5 |
| 15 | Uncharacterized protein | T1ZEQ3 | SIR_0983 | 13 | 7 | 4 | 5 |
| 16 | Foldase protein PrsA | T1ZG93 | prsA | 8 | 7 | 4 | 7 |
| 17 | 30S ribosomal protein S2 | T1ZC88 | rpsB | 2 | 15 | 5 | 3 |
| 18 | Aminopeptidase | T1ZEN3 | pepC | 0 | 21 | 0 | 6 |
| 19 | Pullulanase, type I | T1ZEI5 | pulA | 11 | 5 | 7 | 5 |
| 20 | Elongation factor Tu | T1ZEN1 | tuf | 4 | 9 | 4 | 5 |
| 21 | Protein RecA | T1ZFX5 | recA | 6 | 15 | 0 | 4 |
| 22 | Ribosomal RNA small subunit methyltransferase H | T1ZGX5 | mraW | 6 | 13 | 5 | 3 |
| 23 | Putative rhamnosyltransferase RgpA | T1ZEF3 | rgpA | 0 | 9 | 7 | 7 |
| 24 | 30S ribosomal protein S5 | T1ZGK0 | rpsE | 4 | 8 | 5 | 7 |
| 25 | Oxidoreductase | T1ZD40 | SIR_0796 | 5 | 4 | 8 | 2 |
| 26 | Translation initiation factor IF-3 | T1ZDK2 | infC | 4 | 8 | 0 | 4 |
| 27 | Uracil phosphoribosyltransferase | T1ZGI0 | upp | 3 | 9 | 2 | 4 |
| 28 | Putative lipoprotein | T1ZEA9 | SIR_0850 | 4 | 6 | 6 | 2 |
| 29 | DNA-directed RNA polymerase subunit beta' | T1ZFL9 | rpoC | 4 | 0 | 5 | 5 |
| 30 | Response regulator | T1ZH01 | comE | 4 | 11 | 2 | 0 |
| 31 | Beta-N-acetylhexosaminidase | T1ZED9 | lacZ | 4 | 5 | 2 | 2 |
| 32 | Beta-N-acetylhexosaminidase | T1ZED9-DECOY |  | 6 | 0 | 4 | 2 |
| 33 | L-lactate dehydrogenase | T1ZEP5 | ldh | 3 | 7 | 4 | 4 |
| 34 | 50S ribosomal protein L6 | T1ZGX1 | rplF | 2 | 9 | 0 | 0 |
| 35 | 30S ribosomal protein S12 | T1ZCJ3 | rpsL | 4 | 3 | 4 | 4 |
| 36 | 50S ribosomal protein L4 | T1ZFT4 | rplD | 3 | 2 | 2 | 7 |
| 37 | DNA-binding protein HU | T1ZCZ7 | SIR_0424 | 6 | 2 | 3 | 6 |
| 38 | Uncharacterized protein | T1ZGF5 | SIR_1455 | 2 | 11 | 0 | 0 |
| 39 | Biotin carboxylase | T1ZEV4 | accC | 3 | 7 | 2 | 3 |
| 40 | Uncharacterized protein | T1ZBA4 | SIR_0113 | 4 | 5 | 4 | 0 |
| 41 | Uncharacterized protein | T1ZEG5 | SIR_1274 | 4 | 6 | 0 | 3 |
| 42 | Signal recognition particle protein | T1ZE30 | ffh | 5 | 7 | 0 | 0 |
| 43 | Uncharacterized protein | T1ZGC6 | SIR_1156 | 2 | 5 | 3 | 5 |
| 44 | Mannosyl-glycoprotein endo-beta-N-acetylglucosaminidase | T1ZEU1 | SIR_1072 | 3 | 6 | 2 | 3 |
| 45 | DNA-directed RNA polymerase subunit beta | T1ZGS7 | rpoB | 2 | 5 | 4 | 2 |
| 46 | Surface antigen | T1ZCQ5 | SIR_0054 | 2 | 6 | 5 | 2 |
| 47 | Putative adhesion protein | T1ZDS8 | fszD | 5 | 5 | 0 | 2 |
| 48 | Hyaluronate lyase | T1ZG27 | SIR_1547 | 3 | 6 | 3 | 2 |
| 49 | Putative collagen adhesin | T1ZHC4 | SIR_1805 | 3 | 3 | 5 | 2 |
| 50 | C5a peptidase | T1ZFN2 | SIR_1402 | 3 | 5 | 2 | 2 |
| 51 | Chaperone protein DnaK | T1ZF47 | dnaK | 2 | 5 | 4 | 0 |
| 52 | Putative cell-surface antigen I/II | T1ZHQ3 | SIR_1675 | 0 | 2 | 4 | 0 |
| 53 | Putative glycosyl transferase | T1ZFQ8 | SIR_0933 | 4 | 5 | 0 | 2 |
| 54 | Uncharacterized protein | T1ZFV5 | SIR_1477 | 4 | 3 | 0 | 3 |
| 55 | Formate acetyltransferase | T1ZD63 | pfl | 2 | 5 | 0 | 3 |
| 56 | Lysozyme | T1ZF98 | SIR_1025 | 7 | 2 | 0 | 2 |
| 57 | Translation initiation factor IF-2 | T1ZGZ1 | infB | 0 | 3 | 5 | 2 |
| 58 | Putative alkaline amylopullulanase | T1ZGL9 | pulA2 | 0 | 5 | 4 | 4 |
| 59 | Putative stress protein | T1ZDD5 | SIR_0040 | 0 | 3 | 3 | 0 |
| 60 | 3-oxoacyl-[acyl-carrier-protein] synthase 2 | T1ZH25 | fabF | 2 | 7 | 0 | 2 |
| 61 | DNA polymerase III PolC-type | T1ZGA0 | polC | 3 | 5 | 2 | 0 |
| 62 | Uncharacterized protein | T1ZGR7-DECOY |  | 2 | 0 | 4 | 0 |
| 63 | Glutamine synthetase I alpha | T1ZGF6 | glnA | 0 | 4 | 2 | 5 |
| 64 | ATP-dependent zinc metalloprotease FtsH | T1ZDD0 | ftsH | 3 | 2 | 0 | 4 |
| 65 | LysM domain-containing protein | T1ZHJ6 | SIR_1880 | 4 | 4 | 0 | 0 |
| 66 | Uncharacterized protein | T1ZEQ3-DECOY |  | 0 | 0 | 3 | 0 |
| 67 | Putative phosphoribosylformylglycinamidine synthase | T1ZB52 | purL | 0 | 7 | 2 | 0 |
| 68 | Glyceraldehyde-3-phosphate dehydrogenase | T1ZCF7 | gap | 0 | 7 | 0 | 0 |
| 69 | Pyruvate formate lyase | T1ZF15 | SIR_1079 | 2 | 5 | 0 | 0 |
| 70 | Transcription-repair-coupling factor | T1ZC42 | trcF | 0 | 0 | 0 | 3 |
| 71 | Chromosome partition protein Smc | T1ZDG1 | smc | 0 | 4 | 2 | 0 |
| 72 | Cell division ATP-binding protein FtsE | T1ZF37 | ftsE | 0 | 2 | 2 | 5 |
| 73 | 50S ribosomal protein L18 | T1ZFR9 | rplR | 2 | 3 | 0 | 0 |
| 74 | Alkyl hydroperoxide reductase subunit F | T1ZGT3 | ahpF | 2 | 4 | 0 | 0 |
| 75 | Uncharacterized protein | T1ZCV9-DECOY |  | 0 | 0 | 4 | 0 |
| 76 | Valine--tRNA ligase | T1ZFZ4 | valS | 0 | 4 | 2 | 3 |
| 77 | Phosphoglycerate kinase | T1ZGG2 | pgk | 0 | 2 | 0 | 0 |
| 78 | Beta-N-acetylhexosaminidase | T1ZET7 | SIR_1067 | 2 | 3 | 2 | 0 |
| 79 | 30S ribosomal protein S10 | T1ZH81 | rpsJ | 2 | 3 | 0 | 3 |
| 80 | GRAM_POS_ANCHORING domain-containing protein | T1ZEJ8 | SIR_0758 | 3 | 0 | 0 | 2 |
| 81 | Elongation factor G | T1ZDS4 | fusA | 2 | 3 | 0 | 3 |
| 82 | Lysine--tRNA ligase | T1ZFC7 | lysS | 3 | 2 | 0 | 0 |
| 83 | Putative recombinase | T1ZEH5 | SIR_0971 | 4 | 2 | 0 | 2 |
| 84 | ABC transporter, substrate-binding protein | T1ZD17 | msmE | 0 | 3 | 0 | 0 |
| 85 | Putative conjugal transfer protein | T1ZFW8 | SIR_0990 | 2 | 6 | 0 | 3 |
| 86 | Pyruvate kinase | T1ZEN4 | pyk | 0 | 8 | 0 | 0 |
| 87 | Uncharacterized protein | T1ZCC8 | SIR_0176 | 2 | 3 | 0 | 0 |
| 88 | DD-transpeptidase | T1ZBB4 | SIR_0124 | 3 | 3 | 2 | 0 |
| 89 | DUF4366 domain-containing protein | T1ZDN0 | SIR_0987 | 0 | 2 | 3 | 0 |
| 90 | Type I restriction enzyme R Protein | T1ZDQ2 | hsdR | 3 | 4 | 2 | 0 |
| 91 | Putative DNA-entry endonuclease | T1ZDT4 | endA | 2 | 3 | 0 | 0 |
| 92 | Putative penicillin binding protein 2B | T1ZDK4 | pbp2b | 4 | 0 | 0 | 3 |
| 93 | Threonine--tRNA ligase | T1ZFI3 | thrS | 3 | 3 | 0 | 0 |
| 94 | DUF4832 domain-containing protein | T1ZGK3 | SIR_1591 | 0 | 4 | 4 | 0 |
| 95 | Conjugal transfer protein | T1ZEL6 | SIR_1329 | 4 | 0 | 0 | 0 |
| 96 | Uncharacterized protein | T1ZDP3 | SIR_0613 | 0 | 3 | 6 | 0 |
| 97 | Phosphoenolpyruvate-protein phosphotransferase | T1ZG24 | ptsI | 2 | 2 | 0 | 2 |
| 98 | Isopentenyl-diphosphate delta-isomerase | T1ZD59 | fni | 2 | 5 | 0 | 0 |
| 99 | Peptidyl-prolyl cis-trans isomerase | T1ZE70 | ppiA | 0 | 8 | 0 | 0 |
| 100 | Histidine triad protein | T1ZEB0 | SIR_0654 | 0 | 0 | 2 | 2 |
