## Supplemental Table S2 for "Dual targeting factors are required for LXG toxin export by the bacterial type VIIb secretion system"

**Table S2A. Accession codes and sequence information for LapD2 homologs identified with three iterations of JackHMMER.**

| **Entry** | **Protein names** | **Gene names** | **Organism** | **Length** |
| --- | --- | --- | --- | --- |
| F0ISI0 | Uncharacterized protein | HMPREF9384_0792 | Streptococcus sanguinis SK160 | 120 |
| A0A829IC82 | Uncharacterized protein | SAG0014_09635 | Streptococcus agalactiae FSL S3-586 | 120 |
| A0A427Z096 | Uncharacterized protein | D8894_04900 | Streptococcus oralis | 120 |
| F8DHG2 | Uncharacterized protein | HMPREF0833_11762 | Streptococcus parasanguinis ATCC 15912 | 118 |
| A0A8B1YUD9 | DUF3958 family protein | J4854_01605 | Streptococcus lactarius | 118 |
| A0A178KGP4 | Uncharacterized protein | A3Q39_01935 | Streptococcus sp. CCUG 49591 | 118 |
| A0A1X1IMY3 | Uncharacterized protein | B7710_01130 | Streptococcus oralis subsp. oralis | 120 |
| A0A3R9HBG1 | Uncharacterized protein | D8875_04300 | Streptococcus sanguinis | 120 |
| A3CR32 | Uncharacterized protein | SSA_2275 | Streptococcus sanguinis (strain SK36) | 121 |
| A0A178KI83 | Uncharacterized protein | A3Q39_01965 | Streptococcus sp. CCUG 49591 | 124 |
| S7XHS7 | Uncharacterized protein | M059_05495 | Streptococcus mitis 18/56 | 124 |
| A0A139P9I6 | Uncharacterized protein | SORDD16_01672 | Streptococcus oralis | 121 |
| A0A3R9JF83 | Uncharacterized protein | D8839_01325 | Streptococcus mitis | 118 |
| A0A428A3Y3 | Uncharacterized protein | D8883_04735 | Streptococcus sanguinis | 120 |
| A0A427ZT62 | Uncharacterized protein | D8886_05325 | Streptococcus sanguinis | 120 |
| A0A5A7ZT25 | Uncharacterized protein | FKX92_00600 | Streptococcus sanguinis | 129 |
| A0A7H8V963 | Uncharacterized protein | FFV08_11455 | Streptococcus sanguinis | 120 |
| A0A123VUG4 | FKBP_N domain-containing protein | ERS132372_01528 ERS132399_02391 | Streptococcus suis | 128 |
| A0A428A688 | Uncharacterized protein | D8879_11740 | Streptococcus sanguinis | 120 |
| A0A1F0ZSH8 | Uncharacterized protein | HMPREF2917_09360 | Streptococcus sp. HMSC061E03 | 118 |
| F3UNP6 | Uncharacterized protein | HMPREF9389_0454 | Streptococcus sanguinis SK355 | 121 |
| A0A1X1JWY6 | Uncharacterized protein | B7700_09660 | Streptococcus mitis | 118 |
| A0A345VJJ3 | Uncharacterized protein | Sp14A_09740 | Streptococcus pluranimalium | 129 |
| A0A8B4IQ53 | Uncharacterized protein | NCTC3858_00393 | Streptococcus uberis | 122 |
| A0A0F5MM48 | Uncharacterized protein | RN86_02675 | Streptococcus gordonii | 132 |
| A0A0F2CF76 | Uncharacterized protein | TZ86_01640 UA00_00089 | Streptococcus gordonii | 119 |
| A0A2X3XZG6 | Uncharacterized protein | NCTC12278_01112 | Streptococcus ferus | 131 |
| A0A0E1EH98 | Uncharacterized protein | AX245_04160 C4618_11680 C6N07_05900 RDF_1029 | Streptococcus agalactiae | 118 |
| A0A4T2H8W2 | Uncharacterized protein | FAJ36_02910 | Streptococcus suis | 123 |
| A0A1V0H1D1 | Uncharacterized protein | A6J85_03500 | Streptococcus gordonii | 118 |
| A0A7H8UYG8 | Energy transducer TonB | FDP16_01525 | Streptococcus sanguinis | 120 |
| A0A1E5GHA5 | Uncharacterized protein | BCR21_07310 | Enterococcus ureasiticus | 126 |
| A0A1X1J4E5 | Uncharacterized protein | B7708_00960 | Streptococcus oralis subsp. dentisani | 124 |
| A0A7Z0VFP3 | Uncharacterized protein | TH70_0121 | Streptococcus agalactiae | 123 |
| A0A1E5HGJ1 | Uncharacterized protein | BCR24_01620 | Enterococcus ureilyticus | 118 |
| A0A4P7WQS8 | Uncharacterized protein | E8M06_09955 E8M06_09985 | Streptococcus suis | 123 |
| A0A0U2NRK3 | Uncharacterized protein | ATZ35_10685 | Enterococcus rotai | 118 |
| E6KIR2 | Uncharacterized protein | HMPREF8578_0127 | Streptococcus oralis ATCC 49296 | 120 |
| A0A4R5G734 | Uncharacterized protein | E0E04_02155 | Streptococcus vicugnae | 134 |
| A0A6I3PB65 | Uncharacterized protein | GMC80_04755 GMC84_06710 | Streptococcus parasanguinis | 118 |
| A0A7X2UEL6 | Uncharacterized protein | NCTC3858_01463 | Streptococcus uberis | 126 |
| E6KIP9 | Uncharacterized protein | HMPREF8578_0114 | Streptococcus oralis ATCC 49296 | 118 |
| A0A540UNN3 | FKBP_N domain-containing protein | FH692_10965 | Streptococcus suis | 128 |
| A0A3L8GE13 | Uncharacterized protein | DIY07_08810 | Streptococcus iniae (Streptococcus shiloi) | 125 |
| A0A7H9FG12 | Uncharacterized protein | HRE59_00315 | Streptococcus oralis subsp. oralis | 118 |
| A0A372KJ05 | Uncharacterized protein | DDV21_010945 DDV23_10765 | Streptococcus chenjunshii | 131 |
| A0A0J6KU02 | Uncharacterized protein | VK90_24155 | Bacillus sp. LK2 | 116 |
| A0A427Z4E3 | Energy transducer TonB | D8889_08515 FKX92_06260 | Streptococcus sanguinis | 120 |
| A0A0F5MJX1 | Uncharacterized protein | RN86_02700 | Streptococcus gordonii | 118 |
| A0A7X2UQ75 | Uncharacterized protein | NCTC3858_01475 | Streptococcus uberis | 126 |
| A0A0S3K6Z3 | Uncharacterized protein | ATZ33_01285 | Enterococcus silesiacus | 118 |
| A0A242AUF8 | Uncharacterized protein | A5821_000622 | Enterococcus sp. 7F3_DIV0205 | 120 |
| A0A242H4J5 | Uncharacterized protein | A5866_002132 | Enterococcus sp. 12C11_DIV0727 | 118 |
| A0A242CWU2 | Uncharacterized protein | A5875_003888 | Enterococcus sp. 3H8_DIV0648 | 119 |
| F0IN33 | HD domain protein | HMPREF9383_1536 | Streptococcus sanguinis SK150 | 119 |
| A0A427ZN60 | Uncharacterized protein | D8886_09175 | Streptococcus sanguinis | 120 |
| A0A081QRU4 | Cell-cycle control medial ring component family protein | SK578_0511 | Streptococcus mitis | 124 |
| A0A242ATT0 | Uncharacterized protein | A5821_000410 | Enterococcus sp. 7F3_DIV0205 | 119 |
| A0A3R9J4D5 | Uncharacterized protein | D8860_09785 | Streptococcus oralis | 118 |
| A0A2X3VDB5 | Uncharacterized protein | NCTC11085_00303 | Streptococcus sanguinis | 120 |
| A0A1X1HW15 | Uncharacterized protein | B7714_09145 | Streptococcus oralis subsp. oralis | 120 |
| A0A0Z8JBB1 | Uncharacterized protein | ERS132440_00897 | Streptococcus suis | 123 |
| A0A242GZP3 | Uncharacterized protein | A5866_000650 | Enterococcus sp. 12C11_DIV0727 | 114 |
| A0A841YH39 | DUF3958 family protein | HB844_13135 | Listeria fleischmannii | 118 |
| A0A1E5GX96 | Uncharacterized protein | BCR23_04630 | Enterococcus quebecensis | 115 |
| R2T5H1 | Uncharacterized protein | UAY_00975 | Enterococcus moraviensis ATCC BAA-383 | 118 |
| A0A7H8V9W6 | Uncharacterized protein | FFV08_11490 | Streptococcus sanguinis | 120 |
| F0FHF6 | Uncharacterized protein | HMPREF9388_2139 | Streptococcus sanguinis SK353 | 121 |
| A0A1X1IPR0 | Uncharacterized protein | B7710_00060 | Streptococcus oralis subsp. oralis | 118 |
| A0A428G5R6 | Uncharacterized protein | D8801_04900 | Streptococcus oralis | 124 |
| A0A0N0KTL2 | Uncharacterized protein | AEQ18_02380 | Enterococcus sp. RIT-PI-f | 116 |
| A0A200JBQ9 | Uncharacterized protein | A5889_000138 | Enterococcus sp. 9D6_DIV0238 | 117 |
| A0A7D4GRI0 | Uncharacterized protein | FOC63_06870 | Streptococcus gallolyticus | 134 |
| A0A4T2GM54 | Uncharacterized protein | FAJ39_07710 | Streptococcus suis | 128 |
| A0A242LA88 | Uncharacterized protein | A5881_003618 | Enterococcus termitis | 118 |
| A0A380IM03 | Uncharacterized protein | NCTC6175_01411 | Streptococcus agalactiae | 120 |
| A0A4V6U7E4 | Uncharacterized protein | FAJ36_02880 | Streptococcus suis | 128 |
| A0A3R9HGP9 | Uncharacterized protein | D8887_07705 | Streptococcus sanguinis | 113 |
| A0A7Z7QUJ7 | Uncharacterized protein | NCTC8183_01312 | Streptococcus agalactiae | 133 |
| A0A139NND7 | Uncharacterized protein | STRDD11_02626 | Streptococcus sp. DD11 | 120 |
| A0A2L0D3F4 | Uncharacterized protein | C0J00_04050 | Streptococcus pluranimalium | 131 |
| R2T9D5 | Uncharacterized protein | UAY_02590 | Enterococcus moraviensis ATCC BAA-383 | 117 |
| A0A0Z8HRE2 | Uncharacterized protein | ERS132406_02094 | Streptococcus suis | 123 |
| A0A0B7GNC7 | Uncharacterized protein | SSV_1920 | Streptococcus sanguinis | 120 |
| F9LWN3 | Uncharacterized protein | HMPREF9965_0736 | Streptococcus mitis bv. 2 str. SK95 | 118 |
| A0A1X1JX79 | Uncharacterized protein | B7700_09690 | Streptococcus mitis | 124 |
| A0A1E5H5L3 | Uncharacterized protein | BCR24_09885 | Enterococcus ureilyticus | 122 |
| F0IBB8 | Uncharacterized protein | HMPREF9382_2056 | Streptococcus sanguinis SK115 | 120 |
| R3W643 | Uncharacterized protein | UC3_02024 | Enterococcus phoeniculicola ATCC BAA-412 | 116 |
| A0A428IHC8 | Uncharacterized protein | D8844_06490 | Streptococcus oralis | 120 |
| A0A2W4BKR6 | Uncharacterized protein | CI088_09485 | Enterococcus plantarum | 115 |
| A0A3R9H620 | Uncharacterized protein | D8879_10595 | Streptococcus sanguinis | 120 |
| A0A1E5GJU3 | Uncharacterized protein | BCR25_08215 | Enterococcus termitis | 115 |
| A0A0Z8I4W5 | Uncharacterized protein | ERS132410_02192 | Streptococcus suis | 123 |

**Table S2B. Accession codes and sequence information for LapC1 homologs identified with one iteration of JackHMMER.**

| **Entry** | **Protein names** | **Gene names** | **Organism** | **Length** |
| --- | --- | --- | --- | --- |
| T1ZH75 | Uncharacterized protein | SIR_1491 | Streptococcus intermedius B196 | 91 |
| A0A0E2IQB7 | Uncharacterized protein | HMPREF1654_01870 | Streptococcus intermedius ATCC 27335 | 91 |
| A0A139R5L5 | TIGR04197 family type VII secretion effector | FOC63_00900 SGADD02_00470 SGADD03_00389 | Streptococcus gallolyticus | 93 |
| A0A1S5WDW5 | Uncharacterized protein | BTR42_08900 | Streptococcus gallolyticus subsp. gallolyticus DSM 16831 | 93 |
| A0A1I7GQI7 | Type VII secretion effector, SACOL2603 family | SAMN05660328_102271 | Streptococcus gallolyticus | 93 |
| F5WVX6 | Uncharacterized protein | SGGB_1575 | Streptococcus gallolyticus ATCC 43143 | 93 |
| E8K2Z3 | Uncharacterized protein | HMPREF9423_1856 | Streptococcus infantis ATCC 700779 | 92 |
| A0A1H8Z4E7 | Type VII secretion effector, SACOL2603 family | SAMN05216346_101162 | Streptococcus equinus (Streptococcus bovis) | 90 |
| A0A139QYV5 | Uncharacterized protein | SGADD02_00817 SGADD03_01202 | Streptococcus gallolyticus | 90 |
| F9LY30 | Uncharacterized protein | HMPREF9965_1762 | Streptococcus mitis bv. 2 str. SK95 | 92 |
| A0A1C3SMV2 | Uncharacterized protein | SMA679_0761 | Streptococcus macedonicus | 90 |
| A0A3R9HJH6 | Uncharacterized protein | D8863_08620 | Streptococcus oralis | 92 |
| A0A1H0MTA7 | Type VII secretion effector, SACOL2603 family | SAMN05216347_102469 | Streptococcus equinus (Streptococcus bovis) | 90 |
| A0A371QFB0 | TIGR04197 family type VII secretion effector | DXN33_01140 | Streptococcus sp. NM | 92 |
| A0A3R9QBN4 | Uncharacterized protein | D8786_05750 D8855_04310 | Streptococcus mitis | 92 |
| A0A1F0BUA5 | Type VII secretion protein | HMPREF2613_07245 | Streptococcus sp. HMSC070B10 | 92 |
| A0A501PB50 | TIGR04197 family type VII secretion effector | FJN11_06485 | Streptococcus symci | 92 |
| A0A3R9HQE0 | TIGR04197 family type VII secretion effector | D8789_07065 D8849_09150 D8865_10365 JJN14_03035 | Streptococcus mitis | 92 |
| A0A1E9GAV6 | Type VII secretion protein | HMPREF2766_03755 | Streptococcus sp. HMSC076C08 | 92 |
| A0A2G3NUY4 | TIGR04197 family type VII secretion effector | CS009_05415 CS010_03220 | Streptococcus macedonicus | 90 |
| A0A7D4GS34 | TIGR04197 family type VII secretion effector | FOC63_08560 | Streptococcus gallolyticus | 90 |
| A0A1S5WBI2 | Uncharacterized protein | BTR42_04595 | Streptococcus gallolyticus subsp. gallolyticus DSM 16831 | 90 |
| A0A1B1ID96 | Type VII secretion protein | AXF18_01730 | Streptococcus sp. oral taxon 064 | 92 |
| A0A2I1UMC7 | TIGR04197 family type VII secretion effector | CYK17_09995 | Streptococcus oralis subsp. dentisani | 92 |
| A0A1S0ZA19 | Type VII secretion protein | A7T00_33115 | Salmonella enterica subsp. enterica serovar Saintpaul | 92 |
| A0A380K862 | Type VII secretion effector | NCTC13767_01892 | Streptococcus gallolyticus | 90 |
| A0A1H6SD36 | Type VII secretion effector, SACOL2603 family | SAMN05216460_1192 | Streptococcus sp. 45 | 90 |
| A0A3R9FX19 | Uncharacterized protein | D8894_04980 | Streptococcus oralis | 92 |
| A0A1X1J9V7 | Type VII secretion effector | B7705_06215 | Streptococcus oralis subsp. dentisani | 92 |
| A0A428DJZ0 | Uncharacterized protein | D8847_09950 | Streptococcus mitis | 92 |
| A0A3R9J234 | Uncharacterized protein | D8847_09775 | Streptococcus mitis | 92 |
| F5X0A7 | Uncharacterized protein | SGGB_0839 | Streptococcus gallolyticus ATCC 43143 | 90 |
| A0A139PV09 | Uncharacterized protein | SORDD27_01490 | Streptococcus oralis | 92 |
| I0Q5G0 | Type VII secretion effector, TIGR04197 family | HMPREF1115_1692 | Streptococcus oralis SK610 | 92 |
| A0A1I7FJ84 | Type VII secretion effector, SACOL2603 family | SAMN05660328_101420 | Streptococcus gallolyticus | 90 |
| A0A239RBG6 | Type VII secretion effector, SACOL2603 family | SAMN05216470_0920 | Streptococcus equinus (Streptococcus bovis) | 90 |
| A0A081QNZ9 | Uncharacterized protein | SK578_0768 SMIM3I_00648 SMIM3IV_00595 | Streptococcus mitis | 92 |
| A0A231VWK6 | TIGR04197 family type VII secretion effector | CBI42_08510 | Streptococcus sp. KR | 92 |
| A0A1F0B683 | Type VII secretion protein | HMPREF2701_04775 | Streptococcus sp. HMSC077D04 | 92 |
| A0A4V0BUI7 | Type VII secretion effector | NCTC5338_01391 | Streptococcus australis | 92 |
| A0A4V6LQ02 | Type VII secretion effector | NCTC10232_01364 | Streptococcus oralis | 92 |
| A0A2X3W4X4 | Type VII secretion effector | NCTC12278_01169 | Streptococcus ferus | 91 |
| A0A1X1INN9 | Type VII secretion effector | B7710_01210 | Streptococcus oralis subsp. oralis | 92 |
| A0A3R9KT57 | Uncharacterized protein | D8788_09675 | Streptococcus mitis | 92 |
| J5H474 | Type VII secretion effector, TIGR04197 family | HMPREF1125_0309 | Streptococcus oralis SK304 | 92 |
| A0A1S1CRP1 | Type VII secretion protein | HMPREF2628_07975 | Streptococcus sp. HMSC063B03 | 92 |
| A0A139QMH4 | Uncharacterized protein | SORDD24_01549 | Streptococcus oralis | 92 |
| A0A1X1H983 | Type VII secretion effector | B7721_02930 | Streptococcus oralis subsp. oralis | 92 |
| A0A428IP91 | Uncharacterized protein | D8846_06225 | Streptococcus oralis | 92 |
| A0A1X1HPW8 | Type VII secretion effector | B7716_01660 | Streptococcus oralis subsp. oralis | 92 |
| E9FIV1 | Uncharacterized protein | HMPREF0849_01627 | Streptococcus sp. C300 | 92 |
| A0A139Q4A6 | Type VII secretion protein | BBP19_06505 SORDD30_01629 | Streptococcus oralis | 92 |
| A0A1X1GSZ3 | Type VII secretion effector | B7712_00855 | Streptococcus oralis subsp. oralis | 92 |
| A0A139M8G1 | Uncharacterized protein | SORDD05_01233 | Streptococcus oralis | 92 |
| A0A139QLX9 | Uncharacterized protein | SORDD24_01677 | Streptococcus oralis | 92 |
| A0A139PVN7 | Uncharacterized protein | D8844_06410 SORDD20_00506 | Streptococcus oralis | 92 |
| A0A1X1HNL4 | Type VII secretion effector | B7718_02130 | Streptococcus oralis subsp. oralis | 92 |
| A0A428HB07 | Uncharacterized protein | D8788_03670 | Streptococcus mitis | 92 |
| G6C8X2 | Uncharacterized protein | HMPREF9184_00751 | Streptococcus sp. oral taxon 058 str. F0407 | 92 |
| A0A4Q2FKS1 | TIGR04197 family type VII secretion effector | DF216_07805 | Streptococcus oralis | 92 |
| J5GN34 | Type VII secretion effector, TIGR04197 family | HMPREF1125_2061 | Streptococcus oralis SK304 | 92 |
| A0A1X0X0B5 | Type VII secretion protein | ATE37_07430 | Streptococcus oralis subsp. tigurinus | 92 |
| A0A428CAR2 | Uncharacterized protein | D8856_09625 | Streptococcus mitis | 92 |
| A0A3R9KGB9 | Uncharacterized protein | D8854_03060 | Streptococcus mitis | 92 |
| A0A1X1I2K2 | Type VII secretion effector | B7714_02825 | Streptococcus oralis subsp. oralis | 92 |
| A0A139NUT4 | Uncharacterized protein | SORDD14_01568 | Streptococcus oralis | 92 |
| A0A4Q2FL97 | TIGR04197 family type VII secretion effector | DF216_07290 | Streptococcus oralis | 92 |
| A0A1X1H062 | Type VII secretion effector | B7722_01935 | Streptococcus oralis subsp. oralis | 92 |
| A0A4R5G4Y4 | TIGR04197 family type VII secretion effector | E0E04_04080 | Streptococcus vicugnae | 91 |
| A0A1X1GBF8 | Type VII secretion effector | B7727_03280 | Streptococcus oralis subsp. tigurinus | 93 |
| A0A139NWT6 | Uncharacterized protein | SORDD15_01377 | Streptococcus oralis | 92 |
| A0A135YLC9 | Uncharacterized protein | HMPREF3205_02308 | Streptococcus pasteurianus | 96 |
| A0A1S5WCM9 | Uncharacterized protein | BTR42_05645 | Streptococcus gallolyticus subsp. gallolyticus DSM 16831 | 91 |
| A0A7D4GHM0 | TIGR04197 family type VII secretion effector | FOC63_09620 | Streptococcus gallolyticus | 91 |
| A0A7D4K0Q8 | TIGR04197 family type VII secretion effector | FOC63_07845 | Streptococcus gallolyticus | 91 |
| A0A1I7F6C6 | Type VII secretion effector, SACOL2603 family | SAMN05660328_101220 | Streptococcus gallolyticus | 91 |
| A0A1I7FC93 | Type VII secretion effector, SACOL2603 family | SAMN05660328_101314 | Streptococcus gallolyticus | 91 |
| A0A139NQ79 | Uncharacterized protein | STRDD11_02464 | Streptococcus sp. DD11 | 89 |
| F3USM5 | Uncharacterized protein | HMPREF9389_1833 | Streptococcus sanguinis SK355 | 90 |
| A0A3R9IAM7 | TIGR04197 family type VII secretion effector | D8887_08455 FFV08_05580 | Streptococcus sanguinis | 90 |
| A0A427ZP46 | Uncharacterized protein | D8886_07895 | Streptococcus sanguinis | 90 |
| A0A3R9NTY4 | Uncharacterized protein | D8879_08845 | Streptococcus sanguinis | 90 |
| G5JR52 | Uncharacterized protein | STRCR_1677 STRCR_1937 | Streptococcus criceti HS-6 | 92 |
| A0A2A5SDM4 | TIGR04197 family type VII secretion effector | FEZ46_05180 RU88_GL002128 | Lactococcus raffinolactis | 102 |
| A0A0F3H405 | Uncharacterized protein | TZ97_00642 | Streptococcus parasanguinis | 89 |
| A0A6N3CT23 | Uncharacterized protein | SPLFYP13_01158 | Streptococcus parasanguinis | 89 |
| F8DGG8 | Uncharacterized protein | HMPREF0833_10386 | Streptococcus parasanguinis ATCC 15912 | 94 |
| A0A359YGK2 | Uncharacterized protein | SPADD19_01110 | Streptococcus parasanguinis | 89 |
| A0A1F1A3X5 | Uncharacterized protein | HMPREF2917_04405 | Streptococcus sp. HMSC061E03 | 89 |
| I1ZLJ5 | Uncharacterized protein | Spaf_0919 | Streptococcus parasanguinis FW213 | 103 |
| A0A4Q5BT34 | TIGR04197 family type VII secretion effector | GMC84_09185 GMC94_02205 | Streptococcus parasanguinis | 89 |
| I2NMG3 | Uncharacterized protein | HMPREF9971_1232 | Streptococcus parasanguinis F0449 | 113 |
| G5JRP1 | Uncharacterized protein | STRCR_2050 | Streptococcus criceti HS-6 | 91 |
| A0A1F0AWW4 | Uncharacterized protein | HMPREF2686_08175 | Streptococcus sp. HMSC057G03 | 89 |
| V8BGZ4 | Uncharacterized protein | HMPREF1195_00404 | Streptococcus parasanguinis CC87K | 89 |
| A0A428B5A9 | Uncharacterized protein | D8866_01720 | Streptococcus parasanguinis | 89 |
| A0A4Q2FH31 | TIGR04197 family type VII secretion effector | DF218_03565 | Streptococcus parasanguinis | 89 |
| A0A6I3PR01 | TIGR04197 family type VII secretion effector | GMC95_02245 | Streptococcus parasanguinis | 94 |
| E8K4F1 | Uncharacterized protein | HMPREF8577_0436 | Streptococcus parasanguinis ATCC 903 | 99 |
| A0A6A0B9J2 | Type VII secretion protein | Hs30E_00170 | Lactococcus hodotermopsidis | 101 |
| E3CE17 | Uncharacterized protein | HMPREF9626_1164 | Streptococcus parasanguinis F0405 | 89 |
| A0A4R5G605 | TIGR04197 family type VII secretion effector | E0E04_02150 | Streptococcus vicugnae | 118 |
| A0A7D4GGP0 | TIGR04197 family type VII secretion effector | FOC63_06865 | Streptococcus gallolyticus | 118 |
| A0A139MU55 | Uncharacterized protein | STRDD04_00268 | Streptococcus sp. DD04 | 97 |
| R2R504 | Type VII secretion effector | UAI_02685 | Enterococcus malodoratus ATCC 43197 | 92 |
| A0A8B1YTF9 | TIGR04197 family type VII secretion effector | J4854_05255 | Streptococcus lactarius | 89 |
| A0A224XFQ7 | Uncharacterized protein | RsY01_1995 | Lactococcus reticulitermitis | 102 |
| A0A0A0DFU7 | Uncharacterized protein | SSIN_0557 | Streptococcus sinensis | 104 |
| A0A242DHL2 | Uncharacterized protein | A5875_002996 | Enterococcus sp. 3H8_DIV0648 | 92 |
| A0A7W1YHC9 | TIGR04197 family type VII secretion effector | HPK16_15390 | Listeria rustica | 91 |
| A0A378MC82 | Type VII secretion effector | NCTC10815_01240 | Listeria grayi (Listeria murrayi) | 92 |
| A0A0S3K6Z8 | Uncharacterized protein | ATZ33_01280 | Enterococcus silesiacus | 95 |
| A0A0U2XFK1 | Uncharacterized protein | ATZ35_10680 | Enterococcus rotai | 95 |
| A0A242H2N4 | Uncharacterized protein | A5866_002133 | Enterococcus sp. 12C11_DIV0727 | 95 |
| A0A1E5KVA8 | Uncharacterized protein | BCR26_15430 | Enterococcus rivorum | 93 |
| R2TRA5 | Type VII secretion effector | UAY_00974 | Enterococcus moraviensis ATCC BAA-383 | 95 |
| D7V0H1 | Uncharacterized protein | HMPREF0556_11749 | Listeria grayi DSM 20601 | 95 |
| K8N1C5 | Uncharacterized protein | HMPREF9186_00129 | Streptococcus sp. F0442 | 89 |
| A0A242L9F8 | Uncharacterized protein | A5881_003619 | Enterococcus termitis | 95 |
| A0A1E5HGJ2 | Uncharacterized protein | BCR24_01625 | Enterococcus ureilyticus | 95 |
| A0A2R7ZZP2 | Uncharacterized protein | CDIMF43_180250 CKN86_07930 | Carnobacterium divergens (Lactobacillus divergens) | 92 |
| A0A830LAN8 | TIGR04197 family type VII secretion effector | CW834_00955 | Listeria monocytogenes | 97 |
| A0A242ATT5 | Uncharacterized protein | A5821_000409 | Enterococcus sp. 7F3_DIV0205 | 95 |
| A0A242CX89 | Uncharacterized protein | A5875_003889 | Enterococcus sp. 3H8_DIV0648 | 95 |
| A0A842EF80 | TIGR04197 family type VII secretion effector | HB895_12440 HCB08_04225 HCB25_04225 HCB35_09535 | Listeria booriae | 96 |
| A0A7X0XEW8 | TIGR04197 family type VII secretion effector | HCI99_14105 HCJ13_00955 | Listeria booriae | 96 |
| A0A5E9H6J9 | Type VII secretion effector | NCTC13772_01143 NCTC13772_02346 | Carnobacterium divergens (Lactobacillus divergens) | 92 |
| A0A0J6L2B8 | Type VII secretion effector | VK90_21625 | Bacillus sp. LK2 | 99 |
| A0A3R9G5C1 | Uncharacterized protein | D8887_07710 | Streptococcus sanguinis | 92 |
| A0A5E9H653 | Type VII secretion effector | NCTC13772_02372 | Carnobacterium divergens (Lactobacillus divergens) | 92 |
| A0A7X0WR26 | TIGR04197 family type VII secretion effector | HB856_08660 HCB51_16600 | Listeria booriae | 96 |
| A0A081QQI2 | Uncharacterized protein | D8845_00760 D8855_02220 D8865_04910 SK578_1302 | Streptococcus mitis | 102 |
| A0A428IXG9 | Uncharacterized protein | D8800_00795 | Streptococcus oralis | 102 |
| R0P931 | Uncharacterized protein | D065_00650 | Streptococcus mitis 13/39 | 102 |
| A0A0B7GL02 | Putative type VII secretion effector | SSV_1220 | Streptococcus sanguinis | 97 |
| A0A1E5KUF8 | Uncharacterized protein | BCR26_04480 | Enterococcus rivorum | 93 |
| A0A7Z8G2U5 | Uncharacterized protein | CKN67_07395 | Carnobacterium divergens (Lactobacillus divergens) | 92 |
| A0A8B5GW87 | Uncharacterized protein | CKN75_08770 | Carnobacterium divergens (Lactobacillus divergens) | 92 |
| A0A0S3KD07 | Uncharacterized protein | ATZ33_12420 | Enterococcus silesiacus | 117 |
| R2QLU5 | Type VII secretion effector | UAY_03088 | Enterococcus moraviensis ATCC BAA-383 | 120 |
| A0A7I0FCU7 | Uncharacterized protein | CKN77_09500 | Carnobacterium divergens (Lactobacillus divergens) | 92 |
| A0A242AQ39 | Uncharacterized protein | A5821_003000 | Enterococcus sp. 7F3_DIV0205 | 120 |
| A0A7X9QZ50 | TIGR04197 family type VII secretion effector | HF881_01535 | Streptococcus sp. WB01_FAA12 | 102 |
| W7C7M5 | Uncharacterized protein | MFLO_05320 | Listeria floridensis FSL S10-1187 | 96 |
| A0A2R8A462 | Uncharacterized protein | CDIMF43_50002 CKN69_02300 CKN86_04715 | Carnobacterium divergens (Lactobacillus divergens) | 92 |
| A0A200JBQ5 | Uncharacterized protein | A5889_000137 | Enterococcus sp. 9D6_DIV0238 | 93 |
| A0A6L6HCB9 | TIGR04197 family type VII secretion effector | GIX45_16890 | Erwinia sp. CPCC 100877 | 93 |
| A0A0J6L0K0 | Type VII secretion effector | VK90_24150 | Bacillus sp. LK2 | 99 |
| F0IBB7 | Uncharacterized protein | HMPREF9382_2055 | Streptococcus sanguinis SK115 | 90 |
| A0A346NBA4 | TIGR04197 family type VII secretion effector | DDV21_004010 DDV21_004700 DDV23_11140 | Streptococcus chenjunshii | 93 |
| A0A4R6ZPZ6 | Type VII secretion effector (TIGR04197 family) | DFP96_102255 | Listeria rocourtiae | 96 |
| A0A842AZ36 | TIGR04197 family type VII secretion effector | HCJ13_15535 | Listeria booriae | 102 |
| A0A2C1R825 | TIGR04197 family type VII secretion effector | CON44_02325 | Bacillus cereus | 99 |
| A0A5F0MN81 | Uncharacterized protein | CKN67_04115 CKN75_04550 | Carnobacterium divergens (Lactobacillus divergens) | 92 |
| A0A2W3Z748 | TIGR04197 family type VII secretion effector | CI088_09490 | Enterococcus plantarum | 96 |
| A0A0N0KSY6 | Type VII secretion effector | AEQ18_02375 | Enterococcus sp. RIT-PI-f | 88 |
| A0A7X0T610 | TIGR04197 family type VII secretion effector | HB853_09795 | Listeria welshimeri | 88 |
| A0AFZ6 | Uncharacterized protein | lwe0510 | Listeria welshimeri serovar 6b ATCC 35897 | 88 |
| C5NV31 | Uncharacterized protein | GEMHA0001_1408 | Gemella haemolysans ATCC 10379 | 97 |
| A0A2L0D3S0 | TIGR04197 family type VII secretion effector | C0J00_04045 | Streptococcus pluranimalium | 119 |
| A0A2X3XH13 | Type VII secretion effector | NCTC11085_01347 | Streptococcus sanguinis | 97 |
| F3UAG5 | Uncharacterized protein | HMPREF9393_0458 | Streptococcus sanguinis SK1056 | 97 |
| A0A7H8UYP6 | TIGR04197 family type VII secretion effector | FDP16_01520 | Streptococcus sanguinis | 90 |
| A0A427Z4K1 | Uncharacterized protein | D8889_08520 | Streptococcus sanguinis | 90 |
| A0A0J6L9L6 | Type VII secretion effector | VK90_07905 | Bacillus sp. LK2 | 99 |
| F0IN34 | Uncharacterized protein | HMPREF9383_1537 | Streptococcus sanguinis SK150 | 104 |
| C5NV27 | Uncharacterized protein | GEMHA0001_1404 | Gemella haemolysans ATCC 10379 | 97 |
| A0A6I3IT94 | TIGR04197 family type VII secretion effector | GGH90_02870 | Streptococcus sp. zg-36 | 86 |
| A0A6I3I620 | TIGR04197 family type VII secretion effector | GGG87_02865 | Streptococcus sp. zg-86 | 86 |
| A0A6I4RB72 | TIGR04197 family type VII secretion effector | GGH11_02895 | Streptococcus sp. zg-70 | 102 |
| A0A1E5GIS0 | Type VII secretion effector | BCR25_08220 | Enterococcus termitis | 96 |
| W7C2R9 | Uncharacterized protein | MFLO_13765 | Listeria floridensis FSL S10-1187 | 96 |
| A0A2C6WMQ9 | TIGR04197 family type VII secretion effector | BTJ66_11860 | Staphylococcus edaphicus | 91 |
| A0A5A7ZNX9 | TIGR04197 family type VII secretion effector | FKX92_06255 | Streptococcus sanguinis | 90 |
| A0A1E5GX99 | Type VII secretion effector | BCR23_04625 | Enterococcus quebecensis | 96 |
| A0A2X3V3S3 | Type VII secretion effector | D8883_04730 NCTC11085_00302 | Streptococcus sanguinis | 90 |
| A0A2N6SD26 | TIGR04197 family type VII secretion effector | CJ218_07575 | Gemella sanguinis | 94 |
| A0A1E5H6D9 | Uncharacterized protein | BCR24_09880 | Enterococcus ureilyticus | 92 |
| A0A7Z7QU85 | Type VII secretion effector | NCTC8183_01311 | Streptococcus agalactiae | 126 |
| F3UDH1 | Uncharacterized protein | HMPREF9393_1578 | Streptococcus sanguinis SK1056 | 90 |
| J4X2D6 | Type VII secretion effector, TIGR04197 family | HMPREF1150_0118 | Streptococcus sp. AS14 | 90 |
| A0A0F5MK39 | Uncharacterized protein | RN86_02680 | Streptococcus gordonii | 116 |
| A0A428AH08 | Uncharacterized protein | D8875_04305 | Streptococcus sanguinis | 90 |
| A0A2I1Z9Q6 | TIGR04197 family type VII secretion effector | CYK23_08645 | Streptococcus salivarius | 90 |
| A0A841YI15 | TIGR04197 family type VII secretion effector | HB844_13830 | Listeria fleischmannii | 97 |
| A0A2N6SD59 | TIGR04197 family type VII secretion effector | CJ218_07595 | Gemella sanguinis | 94 |
| A0A2V3VWP8 | Type VII secretion effector (TIGR04197 family) | DFR56_108171 | Pseudogracilibacillus auburnensis | 88 |
| A0A841YHX4 | TIGR04197 family type VII secretion effector | HB844_13140 | Listeria fleischmannii | 96 |
| A0A7X1CAH5 | TIGR04197 family type VII secretion effector | HCJ38_14380 | Listeria immobilis | 97 |
| A0A1J4HAR3 | Type VII secretion protein | HMPREF3241_05535 | Staphylococcus sp. HMSC34G04 | 91 |
| A0A3D8TTD4 | Uncharacterized protein | UR08_00425 | Listeria kieliensis | 97 |
| F0FPX7 | Uncharacterized protein | HMPREF9392_0404 | Streptococcus sanguinis SK678 | 90 |
| F2CGJ8 | Uncharacterized protein | HMPREF9391_1993 | Streptococcus sanguinis SK408 | 90 |
| F0ISH9 | Uncharacterized protein | HMPREF9384_0791 | Streptococcus sanguinis SK160 | 90 |
| G5JNA7 | Uncharacterized protein | STRCR_0144 | Streptococcus criceti HS-6 | 93 |
| A0A1E5GH45 | Uncharacterized protein | BCR21_07315 | Enterococcus ureasiticus | 92 |
| A0A7I0BHX0 | TIGR04197 family type VII secretion effector | E1N03_11860 | Staphylococcus epidermidis | 91 |
| A0A829M3W1 | Type VII secretion protein | M453_0212855 | Staphylococcus epidermidis CIM40 | 91 |
| R2SNU8 | Type VII secretion effector | UAY_02591 | Enterococcus moraviensis ATCC BAA-383 | 96 |
| A0A0B7GN05 | Putative type VII secretion effector | SSV_1921 | Streptococcus sanguinis | 90 |
| W7B2A2 | Uncharacterized protein (Fragment) | MAQA_04586 | Listeria aquatica FSL S10-1188 | 82 |
| A0A1H9PSA2 | Type VII secretion effector, SACOL2603 family | SAMN04488559_10172 | Isobaculum melis | 94 |
| W7B6K0 | Uncharacterized protein (Fragment) | MAQA_04296 | Listeria aquatica FSL S10-1188 | 83 |
| A0A841YE47 | TIGR04197 family type VII secretion effector | HB844_07260 | Listeria fleischmannii | 90 |
| A0AK27 | Uncharacterized protein | lwe1941 | Listeria welshimeri serovar 6b ATCC 35897 | 97 |
| A0A242AUE1 | Uncharacterized protein | A5821_000621 | Enterococcus sp. 7F3_DIV0205 | 92 |
| A0A7X0Y3T3 | TIGR04197 family type VII secretion effector | HCA69_08900 | Listeria grandensis | 90 |
| A0A7X1C884 | TIGR04197 family type VII secretion effector | HCJ38_03345 | Listeria immobilis | 97 |
| W7B9C0 | Uncharacterized protein | MAQA_15976 | Listeria aquatica FSL S10-1188 | 97 |
| A0A172Q5Q7 | Uncharacterized protein | A0O21_01495 | Streptococcus pantholopis | 90 |
| A0A7X0XCD6 | TIGR04197 family type VII secretion effector | HCI99_06655 | Listeria booriae | 90 |
| A0A239X809 | Type VII secretion effector | SAMEA4504048_01597 | Streptococcus acidominimus | 105 |
| V6Z4V5 | Uncharacterized protein | SAG0136_11275 | Streptococcus agalactiae LMG 14747 | 105 |
| A0A540UVH0 | TIGR04197 family type VII secretion effector | FH692_06345 | Streptococcus suis | 106 |
| W7CDF0 | Uncharacterized protein | MFLO_01075 | Listeria floridensis FSL S10-1187 | 97 |
| A0AKF5 | Uncharacterized protein | lwe2069 | Listeria welshimeri serovar 6b ATCC 35897 | 97 |
| A0A7I0AJ13 | TIGR04197 family type VII secretion effector | E1N03_09545 | Staphylococcus epidermidis | 91 |
| A0A7X1C121 | TIGR04197 family type VII secretion effector | HB856_09015 | Listeria booriae | 96 |
| A0A2K4FCE9 | TIGR04197 family type VII secretion effector | CD039_08645 | Staphylococcus argensis | 91 |
| Q8DZR6 | Uncharacterized protein | SAG1032 | Streptococcus agalactiae serotype V ATCC BAA-611 | 85 |
| A0A1F0CEK0 | Uncharacterized protein | HMPREF2570_04395 | Streptococcus sp. HMSC069D09 | 85 |
| J8J5K7 | Uncharacterized protein | IIO_06123 | Bacillus cereus VD115 | 91 |
| A0A1E5L0N0 | Uncharacterized protein | BCR26_07815 | Enterococcus rivorum | 103 |
| C0MDX1 | Uncharacterized protein | SZO_07980 | Streptococcus equi subsp. zooepidemicus (strain H70) | 104 |
| A0A076Z409 | TIGR04197 family type VII secretion effector (Type VII secretion effector) | C4618_05905 D5F95_10620 DK41_05465 NCTC6175_01412 NCTC8185_02368 | Streptococcus agalactiae | 116 |
| A0A829IEV4 | Uncharacterized protein | SAG0014_09640 | Streptococcus agalactiae FSL S3-586 | 116 |
| Q8E5G5 | Uncharacterized protein | gbs1067 | Streptococcus agalactiae serotype III (strain NEM316) | 116 |
| A0A243G320 | Type VII secretion effector | BK774_26435 | Bacillus thuringiensis | 91 |
| A0A428IGV6 | Uncharacterized protein | D8844_06495 | Streptococcus oralis | 121 |
| A0A2S7RWC9 | TIGR04197 family type VII secretion effector | CUS89_04340 | Enterococcus mundtii | 93 |

**Table S2C. Accession codes and sequence information for LapD1 homologs identified with one iteration of JackHMMER.**

| **Entry** | **Protein names** | **Gene names** | **Organism** | **Length** |
| --- | --- | --- | --- | --- |
| A0A1F0ZSZ0 | Type VII secretion protein | HMPREF2917_09355 | Streptococcus sp. HMSC061E03 | 117 |
| A0A359YHE7 | Uncharacterized protein | SPADD19_01412 | Streptococcus parasanguinis | 117 |
| I1ZK44 | Uncharacterized protein | Spaf_0401 | Streptococcus parasanguinis FW213 | 117 |
| A0A2I1TT29 | TIGR04197 family type VII secretion effector | CYK20_05490 | Streptococcus parasanguinis | 117 |
| A0A6I3PAZ6 | TIGR04197 family type VII secretion effector | GMC80_04760 GMC84_06705 | Streptococcus parasanguinis | 117 |
| A0A1V0H196 | TIGR04197 family type VII secretion effector | A6J85_03505 | Streptococcus gordonii | 118 |
| A0A0F5MM43 | Type VII secretion protein | RN86_02705 | Streptococcus gordonii | 118 |
| S7XKY2 | Type VII secretion protein | M059_05530 | Streptococcus mitis 18/56 | 118 |
| A0A1X1L326 | Type VII secretion effector | B7692_08470 B7696_07565 B7700_09665 | Streptococcus mitis | 118 |
| A0A178KGQ9 | Type VII secretion protein | A3Q39_01930 | Streptococcus sp. CCUG 49591 | 118 |
| A0A414PGR1 | TIGR04197 family type VII secretion effector | DW666_08555 | Streptococcus parasanguinis | 117 |
| F8DHG1 | Uncharacterized protein | HMPREF0833_11761 | Streptococcus parasanguinis ATCC 15912 | 117 |
| A0A3R9LZL4 | Uncharacterized protein | D8803_08265 | Streptococcus oralis | 119 |
| A0A428EFW5 | Uncharacterized protein | D8839_01320 | Streptococcus mitis | 119 |
| E6KIQ0 | Uncharacterized protein | HMPREF8578_0115 | Streptococcus oralis ATCC 49296 | 119 |
| A0A8B1YMV5 | TIGR04197 family type VII secretion effector | J4854_01600 | Streptococcus lactarius | 117 |
| F9LWN2 | Uncharacterized protein | HMPREF9965_0735 | Streptococcus mitis bv. 2 str. SK95 | 119 |
| A0A7H9FG17 | TIGR04197 family type VII secretion effector | HRE59_00320 | Streptococcus oralis subsp. oralis | 119 |
| A0A3R9PR96 | Uncharacterized protein | D8860_09790 | Streptococcus oralis | 119 |
| A0A1X1IPJ3 | Type VII secretion effector | B7710_00065 | Streptococcus oralis subsp. oralis | 119 |
| A0A139PJZ1 | Uncharacterized protein | SORDD21_01112 | Streptococcus oralis | 119 |
| A0A3L8GDQ6 | TIGR04197 family type VII secretion effector | DIY07_08815 | Streptococcus iniae (Streptococcus shiloi) | 116 |
| A0A178KI70 | Type VII secretion protein | A3Q39_01960 | Streptococcus sp. CCUG 49591 | 121 |
| A0A1B1IDA9 | Type VII secretion protein | AXF18_01820 | Streptococcus sp. oral taxon 064 | 117 |
| A0A427ZT45 | Uncharacterized protein | D8882_08140 | Streptococcus sanguinis | 128 |
| A3CR33 | Uncharacterized protein | SSA_2276 | Streptococcus sanguinis SK36 | 128 |
| A0A3R9JBV3 | Uncharacterized protein | D8860_05090 | Streptococcus oralis | 117 |
| K0ZUT2 | Uncharacterized protein | GMD4S_06157 | Streptococcus sp. GMD4S | 117 |
| A0A3R9FWZ7 | Uncharacterized protein | D8894_04895 | Streptococcus oralis | 117 |
| K1A200 | Uncharacterized protein | GMD6S_07863 | Streptococcus sp. GMD6S | 117 |
| E6KIR3 | Uncharacterized protein | HMPREF8578_0128 | Streptococcus oralis ATCC 49296 | 117 |
| A0A1X1IMY5 | Type VII secretion effector | B7710_01125 | Streptococcus oralis subsp. oralis | 117 |
| I0Q2A4 | Type VII secretion effector, TIGR04197 family | HMPREF1115_1417 | Streptococcus oralis SK610 | 117 |
| F3UNP7 | Uncharacterized protein | HMPREF9389_0455 | Streptococcus sanguinis SK355 | 128 |
| A0A1X1HVT4 | Type VII secretion effector | B7714_09150 | Streptococcus oralis subsp. oralis | 117 |
| S7XHE0 | Type VII secretion protein | M059_05500 | Streptococcus mitis 18/56 | 121 |
| A0A1X1KD41 | Type VII secretion effector | B7692_08440 B7696_07595 | Streptococcus mitis | 121 |
| A0A428IGV6 | Uncharacterized protein | D8844_06495 | Streptococcus oralis | 121 |
| E3CF41 | Uncharacterized protein | HMPREF9626_1803 | Streptococcus parasanguinis F0405 | 117 |
| A0A1X1JX30 | Type VII secretion effector | B7700_09695 | Streptococcus mitis | 121 |
| F0FHF7 | Uncharacterized protein | HMPREF9388_2140 | Streptococcus sanguinis SK353 | 128 |
| A0A1X1J482 | Type VII secretion effector | B7708_00965 | Streptococcus oralis subsp. dentisani | 121 |
| A0A3R9KBB5 | Uncharacterized protein | D8801_04895 | Streptococcus oralis | 121 |
| A0A076Z409 | TIGR04197 family type VII secretion effector (Type VII secretion effector) | C4618_05905 D5F95_10620 DK41_05465 NCTC6175_01412 NCTC8185_02368 | Streptococcus agalactiae | 116 |
| Q8E5G5 | Uncharacterized protein | gbs1067 | Streptococcus agalactiae serotype III strain NEM316 | 116 |
| A0A829IEV4 | Uncharacterized protein | SAG0014_09640 | Streptococcus agalactiae FSL S3-586 | 116 |
| A0A0E1EMX7 | TIGR04197 family type VII secretion effector | AX245_04155 C4618_11685 C6N07_05895 RDF_1030 | Streptococcus agalactiae | 111 |
| A0A837KW31 | Uncharacterized protein | WA04_10840 | Streptococcus agalactiae | 116 |
| A0A4R5G605 | TIGR04197 family type VII secretion effector | E0E04_02150 | Streptococcus vicugnae | 118 |
| A0A7D4GGP0 | TIGR04197 family type VII secretion effector | FOC63_06865 | Streptococcus gallolyticus | 118 |
| A0A139N5A4 | Uncharacterized protein | SCRDD08_00137 | Streptococcus cristatus | 121 |
| A0A4T2GKS9 | TIGR04197 family type VII secretion effector | FAJ39_07705 | Streptococcus suis | 109 |
| A0A7X2UFL6 | Type VII secretion effector | NCTC3858_01464 | Streptococcus uberis | 111 |
| A0A7Z0VGH5 | Uncharacterized protein | TH70_0120 | Streptococcus agalactiae | 111 |
| A0A8B4IN87 | Type VII secretion effector | NCTC3858_00392 | Streptococcus uberis | 108 |
| A0A7Z7QU85 | Type VII secretion effector | NCTC8183_01311 | Streptococcus agalactiae | 126 |
| Q8DZR6 | Uncharacterized protein | SAG1032 | Streptococcus agalactiae serotype V ATCC BAA-611 | 85 |
| A0A1F0CEK0 | Uncharacterized protein | HMPREF2570_04395 | Streptococcus sp. HMSC069D09 | 85 |
| A0A1E5KUF8 | Uncharacterized protein | BCR26_04480 | Enterococcus rivorum | 93 |
| A0A3R9NTY4 | Uncharacterized protein | D8879_08845 | Streptococcus sanguinis | 90 |
| A0A1E5KVA8 | Uncharacterized protein | BCR26_15430 | Enterococcus rivorum | 93 |
| A0A0F5MK39 | Uncharacterized protein | RN86_02680 | Streptococcus gordonii | 116 |
| A0A7H8V643 | TIGR04197 family type VII secretion effector | FFV08_03635 | Streptococcus sanguinis | 125 |
| F3USM5 | Uncharacterized protein | HMPREF9389_1833 | Streptococcus sanguinis SK355 | 90 |
| A0A3R9IAM7 | TIGR04197 family type VII secretion effector | D8887_08455 FFV08_05580 | Streptococcus sanguinis | 90 |
| A0A427ZP46 | Uncharacterized protein | D8886_07895 | Streptococcus sanguinis | 90 |
| A0A5A7ZT92 | TIGR04197 family type VII secretion effector | FKX92_00595 | Streptococcus sanguinis | 124 |
| A0A139NQ79 | Uncharacterized protein | STRDD11_02464 | Streptococcus sp. DD11 | 89 |
| A0A540UNN4 | TIGR04197 family type VII secretion effector | FH692_10960 | Streptococcus suis | 134 |
| A0A0Z8X7W8 | Type VII secretion effector | ERS132372_01527 ERS132399_02390 | Streptococcus suis | 111 |
| A0A116LSC7 | TIGR04197 family type VII secretion effector (Type VII secretion effector) | ERS132406_02093 ERS132410_02193 FAJ36_02915 | Streptococcus suis | 108 |
| A0A4P7WT47 | TIGR04197 family type VII secretion effector | E8M06_09960 | Streptococcus suis | 108 |
| A0A0Z8DGM0 | TIGR04197 family type VII secretion effector (Type VII secretion effector) | E8M06_09990 ERS132392_00702 JZY07_10375 | Streptococcus suis | 108 |
| A0A0S3K715 | Type VII secretion effector | ATZ33_01365 | Enterococcus silesiacus | 104 |
| A0A1E5HGI4 | Type VII secretion effector | BCR24_01530 | Enterococcus ureilyticus | 104 |
| A0A0B7GN05 | Putative type VII secretion effector | SSV_1921 | Streptococcus sanguinis | 90 |
| A0A239X809 | Type VII secretion effector | SAMEA4504048_01597 | Streptococcus acidominimus | 105 |
| V6Z4V5 | Uncharacterized protein | SAG0136_11275 | Streptococcus agalactiae LMG 14747 | 105 |
| A0A540UVH0 | TIGR04197 family type VII secretion effector | FH692_06345 | Streptococcus suis | 106 |
| A0A242AX92 | Uncharacterized protein | A5821_001500 | Enterococcus sp. 7F3_DIV0205 | 99 |
| A0A139QYV5 | Uncharacterized protein | SGADD02_00817 SGADD03_01202 | Streptococcus gallolyticus | 90 |
| A0A380K862 | Type VII secretion effector | NCTC13767_01892 | Streptococcus gallolyticus | 90 |
| A0A1E5GK70 | Type VII secretion effector | BCR25_06445 | Enterococcus termitis | 104 |
| F0IN34 | Uncharacterized protein | HMPREF9383_1537 | Streptococcus sanguinis SK150 | 104 |
| A0A2G3NUY4 | TIGR04197 family type VII secretion effector | CS009_05415 CS010_03220 | Streptococcus macedonicus | 90 |
| A0A3R9G5C1 | Uncharacterized protein | D8887_07710 | Streptococcus sanguinis | 92 |
| A0A0A0DFU7 | Uncharacterized protein | SSIN_0557 | Streptococcus sinensis | 104 |
| A0A2L0D3S0 | TIGR04197 family type VII secretion effector | C0J00_04045 | Streptococcus pluranimalium | 119 |
| A0A4T2H474 | TIGR04197 family type VII secretion effector | FAJ36_02885 | Streptococcus suis | 134 |
| F5X0A7 | Uncharacterized protein | SGGB_0839 | Streptococcus gallolyticus ATCC 43143 | 90 |
| A0A7D4GS34 | TIGR04197 family type VII secretion effector | FOC63_08560 | Streptococcus gallolyticus | 90 |
| A0A1S5WBI2 | Uncharacterized protein | BTR42_04595 | Streptococcus gallolyticus subsp. gallolyticus DSM 16831 | 90 |
| A0A359YGK2 | Uncharacterized protein | SPADD19_01110 | Streptococcus parasanguinis | 89 |
| A0A242H2M5 | Uncharacterized protein | A5866_002123 | Enterococcus sp. 12C11_DIV0727 | 103 |
| A0A7H8UYP6 | TIGR04197 family type VII secretion effector | FDP16_01520 | Streptococcus sanguinis | 90 |
| A0A0U2NRL1 | Type VII secretion effector | ATZ35_10775 | Enterococcus rotai | 103 |
| A0A4Q2FH31 | TIGR04197 family type VII secretion effector | DF218_03565 | Streptococcus parasanguinis | 89 |
| A0A1I7FJ84 | Type VII secretion effector, SACOL2603 family | SAMN05660328_101420 | Streptococcus gallolyticus | 90 |
| A0A1E5GE80 | Type VII secretion effector | BCR21_11960 | Enterococcus ureasiticus | 103 |
| A0A428AH08 | Uncharacterized protein | D8875_04305 | Streptococcus sanguinis | 90 |
| A0A2I1Z9Q6 | TIGR04197 family type VII secretion effector | CYK23_08645 | Streptococcus salivarius | 90 |
| F3UDH1 | Uncharacterized protein | HMPREF9393_1578 | Streptococcus sanguinis SK1056 | 90 |
| I2NMG3 | Uncharacterized protein | HMPREF9971_1232 | Streptococcus parasanguinis F0449 | 113 |
| A0A242LA78 | Uncharacterized protein | A5881_003608 | Enterococcus termitis | 104 |
| A0A6I3PR01 | TIGR04197 family type VII secretion effector | GMC95_02245 | Streptococcus parasanguinis | 94 |
| A0A0E2IQB7 | Uncharacterized protein | HMPREF1654_01870 | Streptococcus intermedius ATCC 27335 | 91 |
| R2QN21 | Type VII secretion effector | UAY_02986 | Enterococcus moraviensis ATCC BAA-383 | 98 |
| A0A139MU55 | Uncharacterized protein | STRDD04_00268 | Streptococcus sp. DD04 | 97 |
| A0A6N3CT23 | Uncharacterized protein | SPLFYP13_01158 | Streptococcus parasanguinis | 89 |
| A0A1F1A3X5 | Uncharacterized protein | HMPREF2917_04405 | Streptococcus sp. HMSC061E03 | 89 |
| A0A427Z4K1 | Uncharacterized protein | D8889_08520 | Streptococcus sanguinis | 90 |
| A0A1F0AWW4 | Uncharacterized protein | HMPREF2686_08175 | Streptococcus sp. HMSC057G03 | 89 |
| J4X2D6 | Type VII secretion effector, TIGR04197 family | HMPREF1150_0118 | Streptococcus sp. AS14 | 90 |
| A0A0B7GL02 | Putative type VII secretion effector | SSV_1220 | Streptococcus sanguinis | 97 |
| A0A4Q5BT34 | TIGR04197 family type VII secretion effector | GMC84_09185 GMC94_02205 | Streptococcus parasanguinis | 89 |
| F8DGG8 | Uncharacterized protein | HMPREF0833_10386 | Streptococcus parasanguinis ATCC 15912 | 94 |
| E8K4F1 | Uncharacterized protein | HMPREF8577_0436 | Streptococcus parasanguinis ATCC 903 | 99 |
