## Supplemental Table S3 for "Dual targeting factors are required for LXG toxin export by the bacterial type VIIb secretion system"

**Table S3. Strains used in this study**

| **Organism** | **Genotype** | **Description** | **Reference** |
| --- | --- | --- | --- |
| *S. intermedius* B196 | Wildtype |  | (1) |
|  | ΔSIR_1490::kan^R^ | *lapC2* deletion strain | This study |
|  | ΔSIR_0175::kan^R^ | *essC* deletion strain | (2) |
|  | ΔSIR_1489-1486::kan^R^ | *telC*-*tipC2* deletion strain | (3) |
| *S. intermedius* GC1825 | Wildtype |  |  |
|  | ΔGC1825_00249::kan^R^ | *esxA* deletion strain | This study |
|  | ΔGC1825_00255::kan^R^ | *lapD1* deletion strain | This study |
|  | ΔGC1825_00256::kan^R^ | *lapD2* deletion strain | This study |
| *E. coli* XL-1 Blue | *recA1 endA1 gyrA96 thi-1 hsdR17 supE44 relA1 lac* [*F’ proAB lacI^q^* Z Δ M15 Tn*10* (Tet^R^)] | Cloning strain. | Agilent |
| *E. coli* BL21 (DE3) CodonPlus | F^-^ *ompT gal dcm lon hsdS*_B_(r_B_^-^ m_B_^-^) λ(DE3) pLysS(Cm^R^) | Protein expression strain. | Novagen |
| *E. coli* B834 (DE3) | F^-^ *ompT gal dcm hsdS*_B_(r_B_^-^ m_B_^-^) λ(DE3) *met* | Protein expression methionine auxotroph. | Novagen |
