## Supplemental Table S4 for "Dual targeting factors are required for LXG toxin export by the bacterial type VIIb secretion system"

**Table S4. Plasmids used in this study**

| **Plasmid** | **Relevant features** | **Reference** |
| --- | --- | --- |
| pDL277 | *Streptococcus*-*E. coli* shuttle vector, Spec^R^ | (4) |
| pETDuet-1 | Co-expression vector with *lacI*, T7 promoter, N-terminal His_6_ tag in MCS1, Amp^R^ | Novagen |
| pET29b | Expression vector with *lacI*, T7 promoter, C-terminal His_6_ tag, Kan^R^ | Novagen |
| pSCrhaB2 | Expression vector with *PrhaB*, Tmp^R^ | (5) |
| pPSV39 | Expression vector with *lacI*, *lacUV5* promoter, GmR | (6) |
| pDL277::P96_*lapC1*_VSV-G | *S. intermedius* expression vector for LapC1, C-terminal VSV-G tag | This study |
| pDL277::P96_*lapC2*_VSV-G | *S. intermedius* expression vector for LapC2, C-terminal VSV-G tag | This study |
| pDL277::P96_*telC*_VSV-G | *S. intermedius* expression vector for TelC, C-terminal VSV-G tag | (3) |
| pDL277::P96_lapD1_VSV-G | *S. intermedius* expression vector for LapD1, C-terminal VSV-G tag | This study |
| pDL277::P96_*lapD1*_F77A__VSV-G | *S. intermedius* expression vector for LapD1 with an F77A mutation, C-terminal VSV-G tag | This study |
| pDL277::P96_*lapD1*_D81A__VSV-G | *S. intermedius* expression vector for LapD1 with an D81A mutation, C-terminal VSV-G tag | This study |
| pDL277::P96_*lapD2*_VSV-G | *S. intermedius* expression vector for LapD2, C-terminal VSV-G tag | This study |
| pDL277::P96_*lapD2*_C59S__VSV-G | *S. intermedius* expression vector for LapD2 with an C59S mutation, C-terminal VSV-G tag | This study |
| pDL277::P96_*esxA*_VSV-G | *S. intermedius* expression vector for *esxA* from *Si* GC1825 with a C-terminal VSV-G tag | This study |
| pETDuet-1::*telC*_LXG__His_6_::*lapC1* | *E. coli* co-expression vector for the LXG domain of TelC with LapC1, C-terminal His_6_ on TelC | This study |
| pETDuet-1::*telD*_LXG__His_6_::*lapD1* | *E. coli* co-expression vector for the LXG domain of TelD with LapD1, C-terminal His_6_ on TelD | This study |
| pETDuet-1::*telD*_LXG__His_6_::*lapD1*_F77A_ | *E. coli* co-expression vector for the LXG domain of TelD with LapD1_F77A_, C-terminal His_6_ on TelD | This study |
| pETDuet-1::*telD*_LXG__His_6_::*lapD1*_D81A_ | *E. coli* co-expression vector for the LXG domain of TelD with LapD1_D81A_, C-terminal His_6_ on TelD | This study |
| pET29b::*lapC2* | *E. coli* expression vector for LapC2 | This study |
| pET29b::*lapD2* | *E. coli* expression vector for LapD2 | This study |
| pET29b::*lapD2*_C59S_ | *E. coli* expression vector for LapD2_C59S_ | This study |
| pET29b::*lapD2*_His_6_ | *E. coli* expression vector for LapD2, C-terminal His_6_ tag | This study |
| pET29b::*lapD2*_C59S__His_6_ | *E. coli* expression vector for LapD2_C59S_, C-terminal His_6_ tag | This study |
| pSCrhaB2::*telD* | Rhamnose inducible expression of TelD | This study |
| pPSV39::*tipD* | IPTG inducible expression of TipD | This study |
